# Myeloid progenitor dysregulation fuels immunosuppressive macrophages in tumors

**DOI:** 10.1101/2024.06.24.600383

**Authors:** Samarth Hegde, Bruno Giotti, Brian Y. Soong, Laszlo Halasz, Jessica Le Berichel, Assaf Magen, Benoit Kloeckner, Raphaël Mattiuz, Matthew D. Park, Adam Marks, Meriem Belabed, Pauline Hamon, Theodore Chin, Leanna Troncoso, Juliana J. Lee, Dughan Ahimovic, Michael Bale, Grace Chung, Darwin D’souza, Krista Angeliadis, Travis Dawson, Seunghee Kim-Schulze, Raja M. Flores, Andrew J. Kaufman, Florent Ginhoux, Steven Z. Josefowicz, Sai Ma, Alexander M. Tsankov, Thomas U. Marron, Brian D. Brown, Miriam Merad

**Affiliations:** Marc and Jennifer Lipschultz Precision Immunology Institute, Icahn School of Medicine at Mount Sinai; Department of Immunology and Immunotherapy, Icahn School of Medicine at Mount Sinai; Department of Genetics and Genomic Sciences, Icahn School of Medicine at Mount Sinai; The Human Immune Monitoring Center, Icahn School of Medicine at Mount Sinai; Division of Hematology/Oncology, Icahn School of Medicine at Mount Sinai; Center for Thoracic Oncology, Icahn School of Medicine at Mount Sinai; The Tisch Cancer Institute, Icahn School of Medicine at Mount Sinai, New York, NY, USA; Gustave Roussy Cancer Campus, Villejuif, France; Institut National de la Santé Et de la Recherche Médicale (INSERM) U1015, Equipe Labellisée-Ligue Nationale contre le Cancer, Villejuif, France; Department of Immunology, Harvard Medical School, Boston, MA, USA; Immunology and Microbial Pathogenesis Program, Weill Cornell Graduate School of Medical Sciences; Laboratory of Epigenetics and Immunity, Department of Pathology and Laboratory Medicine, Weill Cornell Medicine, Cornell University, New York, NY, USA

## Abstract

Monocyte–derived macrophages (mo-macs) drive immunosuppression in the tumor microenvironment (TME) and tumor-enhanced myelopoiesis in the bone marrow (BM) fuels these populations. Here, we performed paired transcriptome and chromatin analysis over the continuum of BM myeloid progenitors, circulating monocytes, and tumor-infiltrating mo-macs in mice and in patients with lung cancer to identify myeloid progenitor programs that fuel pro-tumorigenic mo-macs. Analyzing chromatin accessibility and histone mark changes, we show that lung tumors prime accessibility for Nfe2l2 (NRF2) in BM myeloid progenitors as a cytoprotective response to oxidative stress. NRF2 activity is sustained and increased during monocyte differentiation into mo-macs in the lung TME to regulate oxidative stress, in turn promoting metabolic adaptation, resistance to cell death, and contributing to immunosuppressive phenotype. NRF2 genetic deletion and pharmacological inhibition significantly reduced mo-macs’ survival and immunosuppression in the TME, enabling NK and T cell therapeutic antitumor immunity and synergizing with checkpoint blockade strategies. Altogether, our study identifies a targetable epigenetic node of myeloid progenitor dysregulation that sustains immunoregulatory mo-macs in the TME.

## MAIN

A major focus in cancer immunotherapy has been reprogramming monocyte–derived macrophages (mo- macs) in the tumor microenvironment (TME) of solid tumors to reverse immunosuppression and unleash T cell/NK cell responses^1^. While this TME–centric approach has considerable merit, it fails to tackle the ‘wellspring’ of bone marrow (BM) myeloid progenitors seeding monocytes and mo-macs in the TME via tumor-driven myelopoiesis^2–5^. Demand-adapted myelopoietic mobilization during infection/trauma is linked to transcriptomic changes in BM myeloid progenitors enabling their expansion and survival^6–8^, but we have not yet deciphered the exact nature of epigenetic and metabolic changes that occur in myeloid progenitors and mo-mac progeny during tumor-induced myelopoiesis. As our view of systemic tumor– host crosstalk expands, it becomes important to understand if and how complex tumoral cues can ‘pre- condition’ myeloid progenitors in BM by altering their chromatin states, priming gene programs of immunoregulation and undermining anti-tumor responses. Understanding such regulation during chronic tumor inflammation is key to therapeutically targeting the mo-mac replenishment cycle and developing more durable myeloid-targeting therapies. Here, we sought to identify genetic and epigenetic changes that prime myeloid progenitors in tumor-bearing hosts, probe their contribution to tumorigenic immunoregulatory mo-mac plasticity in the TME, and harness this knowledge to redirect TME mo-mac fate towards an anti-tumor phenotype.

### Pathogenic myelopoiesis in lung cancer associates with changes in the chromatin state of BM myeloid progenitors

To initially characterize the impact of lung cancer growth on BM myelopoiesis, we profiled progenitors in BM of naïve and *Kras*^LSL-G12D/+^*; Trp53*^fl/fl^ (KP)^9, 10^ tumor-bearing mice at early (day 7), middle (day 15), and advanced (day 21+) timepoints using multiparametric flow cytometry. We observed a marked increase in hematopoietic stem cells (HSC-LT), granulocytic-monocytic progenitors (GMP) and committed monocyte progenitor (cMoP) by flow cytometry in BM of advanced tumor–bearing mice (**Fig. 1A**). In support of myeloid-biased expansion, BM progenitors from late-stage tumor-bearing mice formed markedly more granulocytic-monocytic colonies (CFU-GM) compared to erythroid colonies (BFU-E) and proliferated more than their naïve counterparts (**Extended Data Fig. 1A**). In addition, we observed an increased mobilization of Ly6C^hi^ monocytes and Ly6G^hi^ neutrophils in circulation tracking with increased tumor burden (**Fig. 1B**). We also observed an expansion of GMP–derived CD157^+^ or CD177^+^ Ly6C^hi^ neutrophil- like monocytes in the BM and peripheral blood of late-stage tumor–bearing mice (**Fig. 1C**). These GMP– derived monocytes have been described to expand in emergency inflammatory conditions and engender oxidative burst-associated degranulation gene programs^11, 12^.

**Fig. 1:**
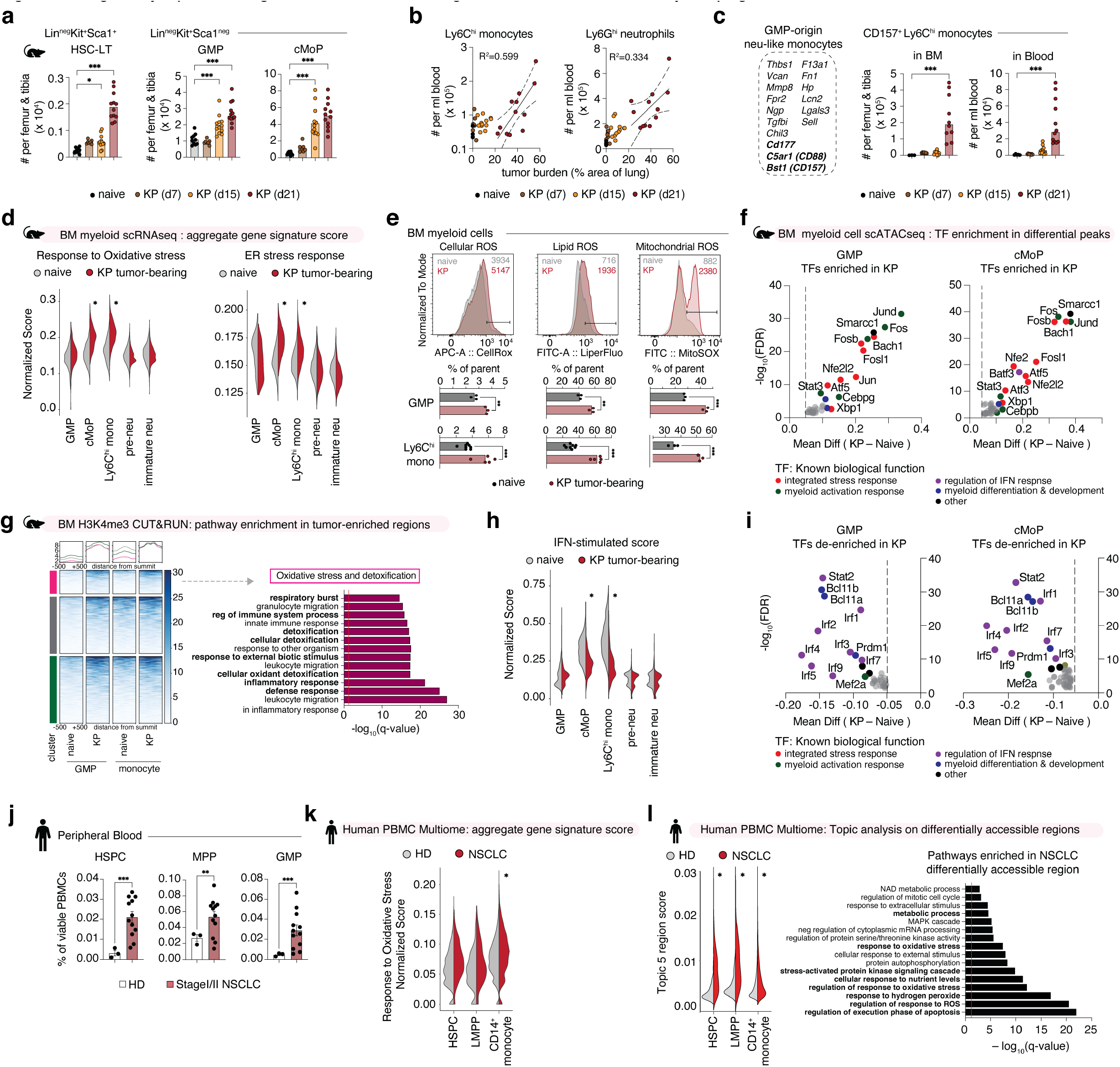
Pathogenic myelopoiesis in cancer associates with changes in the chromatin state of bone marrow myeloid progenitors a. Abundance of long-term hematopoietic stem cells (HSC-LT), granulocytic-monocytic progenitors (GMP), and committed monocytic progenitors (cMoP) in bone marrow (BM) of naïve and KP tumor-bearing mice at different time points. Pooled from two independent experiments. N=7–12 mice per group. Data are individual data points with bar denoting mean. b. Abundance of Ly6C^hi^ monocytes and Ly6G^hi^ neutrophils in blood of naïve and KP tumor-bearing mice at different time points, correlated with tumor burden. Pooled from two independent experiments. N=7–12 mice per group. Data are individual data points with confidence intervals for linear regression. c. Gene markers characterizing GMP-origin neutrophil-like monocytes, and abundance of CD115^+^Ly6C^hi^CD157^+^ monocytes in BM and blood of naïve and KP tumor-bearing mice at different time points. Pooled from two independent experiments. N=3–12 mice per group. Data are individual data points with bar denoting mean. d. Normalized UCell-computed scores for Oxidative stress response and ER stress response gene signatures in select BM cell states from scRNA-seq of naïve and KP tumor-bearing mice. Pooled from 3 mice for one experiment. Data are per-cell distribution violin plot. e. Representative flow cytometry histograms and quantification of CellROX, LiperFluo (lipid peroxidation indicator), and MitoSOX (mitochondrial oxidative burden indicator) in viable BM GMPs and CD115^+^Ly6C^hi^ monocytes from naïve and KP tumor-bearing mice. N=3–7 mice per group. f. Transcription Factor (TF) motifs differentially enriched in KP-tumor bearing mouse BM GMPs (LEFT) and BM cMoPs (RIGHT) compared to naïve mice; ranked by false discovery rate. Dot color indicating major known biological pathways. g. Clusters of H3K4me3 regions enriched in BM GMPs and Ly6C^hi^ monocytes from KP tumor-bearing mice compared to naïve mice (left), with pathways enriched in indicated cluster involving oxidative stress and detoxification (right). Curated terms arranged by adjusted p-value (log q-value). h. Normalized UCell-computed scores for Type I/III Interferon (IFN)-stimulated gene signature in select BM cell states from scRNA-seq of naïve and KP tumor-bearing mice. Pooled from 3 mice for one experiment. Data are per-cell distribution violin plot. i. TF motifs differentially de-enriched in BM GMPs (LEFT) and BM cMoPs (RIGHT) from KP tumor-bearing mice vs naïve mice; ranked by false discovery rate. Dot color indicating major known biological pathways. j. Frequency of hematopoietic stem and progenitor cells (HSPCs), multipotent progenitors (MPP), and granulocytic-monocytic precursors (GMPs) from blood of patients with NSCLC (N=9) and healthy donors (HD, N=3). Pooled from three independent experiments. k. Normalized UCell-computed scores for Oxidative stress response gene signature in indicated PBMC myeloid cells from patients with NSCLC (N=3) and healthy donors (HD, N=2). Data are per-cell distribution violin plot. l. Region scores for topic associated with differentially accessible regions enriched in HSPCs and CD14 monocytes from blood of patients with NSCLC compared to healthy donors (left), with pathways enriched in indicated topic (right). Data are per-cell distribution violin plot (left). *p*-values computed by one-way ANOVA with Dunnett’s multiple comparisons test (A),(C),(D),(H),(K),(L). *p*-values computed by unpaired t-test across conditions (E),(J). *p*-values computed by hypergeometric test with multiple test correction (G),(L). P-value of < 0.05 denoted *; p-values < 0.01 denoted **; p-values < 0.001 denoted ***.

Given this phenotypic shift during myeloid expansion in response to malignancy, we probed how molecular programs are transcriptionally and epigenetically rewired in myeloid progenitors during tumor-induced myelopoiesis. We directed our attention to monocytic lineage progenitors (GMP and cMoP) as they are the main source of suppressive mo-macs in tumors^13, 14^. Experimentally, we utilized scRNA-seq to interrogate myeloid progenitor lineages (Lineage^neg^ CD117^hi^ Sca1^neg^ CD34^+^ CD16/32^+^) sorted from the BM of age-matched naïve and advanced-stage tumor-bearing mice. We could identify distinct granulocytic lineage i.e. GP, pre-neutrophil, immature neutrophil, mature neutrophil^15^, and monocytic lineage i.e. GMP, cMoP, Ly6C^hi^ monocyte, Ly6C^low^ monocyte^16^, delineated by established gene markers (**Extended Data Fig 1B**). Importantly, differentially upregulated genes in tumor-associated GMPs and cMoPs were associated with biological processes involving *response to reactive oxygen species (ROS), hypoxia response, regulation of apoptosis*, and *metabolic regulation of superoxide generation* (**Extended Data Fig 1C**). When we analyzed public datasets of BM HSCs and myeloid progenitors isolated from PyMT breast cancer-bearing mouse models^17, 18^, we found similar enrichment of genes involved in *response to ROS, response to hypoxia,* and *integrated stress response/*endoplasmic reticulum (*ER) stress response* (**Extended Data Fig. 1D**). We orthogonally found tumor-associated BM cMoPs and Ly6C^hi^ monocytes to have higher gene signature scores for oxidative stress response and ER stress response (**Fig. 1D**; gene list in **Supplementary Table 1**). Using flow cytometry, we confirmed that BM GMPs and Ly6C^hi^ monocytes from tumor-bearing mice had increased cellular ROS burden, lipid peroxidation, and mitochondrial oxidative stress (**Fig. 1E**). Thus, across three datasets spanning two different models of cancer, we identified conserved pathways of response to ROS stress and apoptosis regulation upregulated in tumor-associated BM myeloid progenitors.

We next focused on understanding the genomic changes influencing transcription factor (TF) occupancy which drive downstream gene programs during tumor-induced myelopoiesis. To do so, we integrated the scRNA-seq profile of myeloid cells with paired scATAC-seq data from the same experiment, which facilitated an unprecedented granular look at co-regulated gene programs within each myeloid cell state and correlation to chromatin accessibility profiles (**Extended Data Fig. 1E**). We could verify that differentially accessible marker regions for GMPs were enriched in TF motifs such as *Gata2, Tal1, Lyl1*^19, 20^, while cMoPs and monocytes had increased motif enrichment for well-established TFs such as, *Irf4, Cebpb*^21^ and *Spib*^22^ (**Extended Data Fig. 1F**). Having established the transcriptional changes and accessible chromatin regions for each BM myeloid cluster, we sought to identify TF motifs enriched in differentially accessible regions of tumor-associated cell states. In line with our transcriptional results, tumor-associated GMPs and cMoPs had increased motif accessibility for cytoprotective oxidative stress-responsive TFs such as *Nfe2l2, Bach1, Fosl2, Atf3, Atf5, Nfil3, Stat3, and Xbp1*^23^, AP–1 stress-response factors *Jund, Fos,* and granulocytic fate regulators *Cebpd and Cebpa*^15, 16^ (**Fig. 1F**), when compared to naïve counterparts. These observations were further supported by our analysis of H3K4me3 CUT&RUN data on tumor-associated BM GMPs and monocytes; here we observed H3K4me3 signal (denoting poised and active promoter regions) was gained in tumor-associated GMPs and monocytes in genomic regions associated with oxidative stress handling, detoxification, chromatin remodeling, and proliferation (**Fig. 1G** and **Extended Data Fig. 1G**). On the other hand, we observed a distinct reduction of Type-I/III interferon (IFN)-stimulated gene expression in tumor-associated BM cMoPs and pre-monocytes, implying a dampened inflammatory state^24, 25^ distal to the TME (**Fig. 1H**; gene list in **Supplementary Table 1**). We found reduced chromatin accessibility for several IFN pathway TFs including *Irf3, Irf7, Irf5,* and *Stat2* in tumor-associated GMPs and cMoPs (**Fig. 1I**), supporting our transcriptomic findings of IFN hypo- sensitivity. These results collectively suggested that tumor-associated myelopoiesis drives poising and potential activation of pathways in BM GMPs encompassing proliferation-induced ROS stress response^26^, mitochondrial and ER metabolic adaptations^27^, and IFN hypo-responsiveness to prevent chronic IFN-dependent exhaustion^28^.

Subsequently, we quantified mobilized progenitor populations in the blood of patients with early-stage NSCLC and found that CD34^+^CD38^neg^CD90^+^CD49f^+^ HSCs, CD34^+^CD38^neg^CD90^neg^CD49f^neg^ multipotent progenitors (MPPs), and CD34^+^CD38^+^ GMPs were much more abundant in peripheral blood from patients with early-stage NSCLC when compared to healthy donors (**Fig. 1J**). Mirroring our murine transcriptional findings, oxidative stress response gene signature scores were higher in CD14^+^ monocytes from peripheral blood of patients with NSCLC when compared to healthy donors (**Fig. 1K**). We found the gene programs differentially expressed in CD14^+^ monocytes from patients with NSCLC to be downstream of TF regulators such as *NFE2L2, STAT3, PPARG,* and *SMAD4* (**Extended Data Fig. 1H**). Epigenetically, the differentially accessible regions in CD14^+^ monocytes from patients with NSCLC were enriched for genes associated with *metabolic processes, response to ROS and response to oxidative stress* (**Fig. 1L**) downstream of TFs such as *NFE2L2* and *STAT3*. Thus, we found tumor-induced myelopoiesis to drive stress-responsive transcriptomic and epigenetic changes in the mobilized monocytic lineage of patients with NSCLC, recapitulating our findings from murine tumor models. Our data collectively demonstrate that cancer-associated inflammation provokes demand-adapted mobilization of HSCs and multipotent progenitors with myeloid skewing^29–31^, and that this mobilization can impart genetic programs and imprint epigenomic states of BM myeloid progenitors for oxidative stress response and IFN hypo- responsiveness.

### Tumor myelopoiesis fuels mo-macs in TME with sustained cytoprotective stress response

We evaluated if such imprinting of stress-responsive genetic programs in BM myeloid progenitors upon sensing tumor cues can influence the subsequent development of monocytic-macrophages (mo-macs) in the TME. To do so, we traced the fate of transferred GMPs primed in tumor-bearing mice, and found the CD64^+^MERTK^+^ mo-macs derived from tumor-primed GMPs to be more immunoregulatory in the TME, characterized by increased presence of GPNMB^+^CD9^+^ TREM2^hi^ mo-macs^32, 33^ expressing higher PD–L1, increased Arg1^+^ mo-macs^34^, and decreased CD86^+^MHCII^+^ immunostimulatory mo-macs (**Fig. 2A**). In addition, we utilized an *ex vivo* model of tumor conditioning early in bone marrow-derived macrophage (BMDM) differentiation and compared it to tumor exposure later in differentiation. The exposure to tumor inflammatory cues early in the differentiation trajectory resulted in BMDMs with increased Arg1 and PDL1 expression, and decreased MHCII and CD86 (**Fig. 2B**). These results collectively implied that initial exposure of myeloid progenitors to tumor cues impacted their ultimate trajectory in the TME; we therefore proceeded to study the gene regulation of progeny monocytes and mo-macs in the TME in relation to initial changes in BM progenitors.

**Fig. 2:**
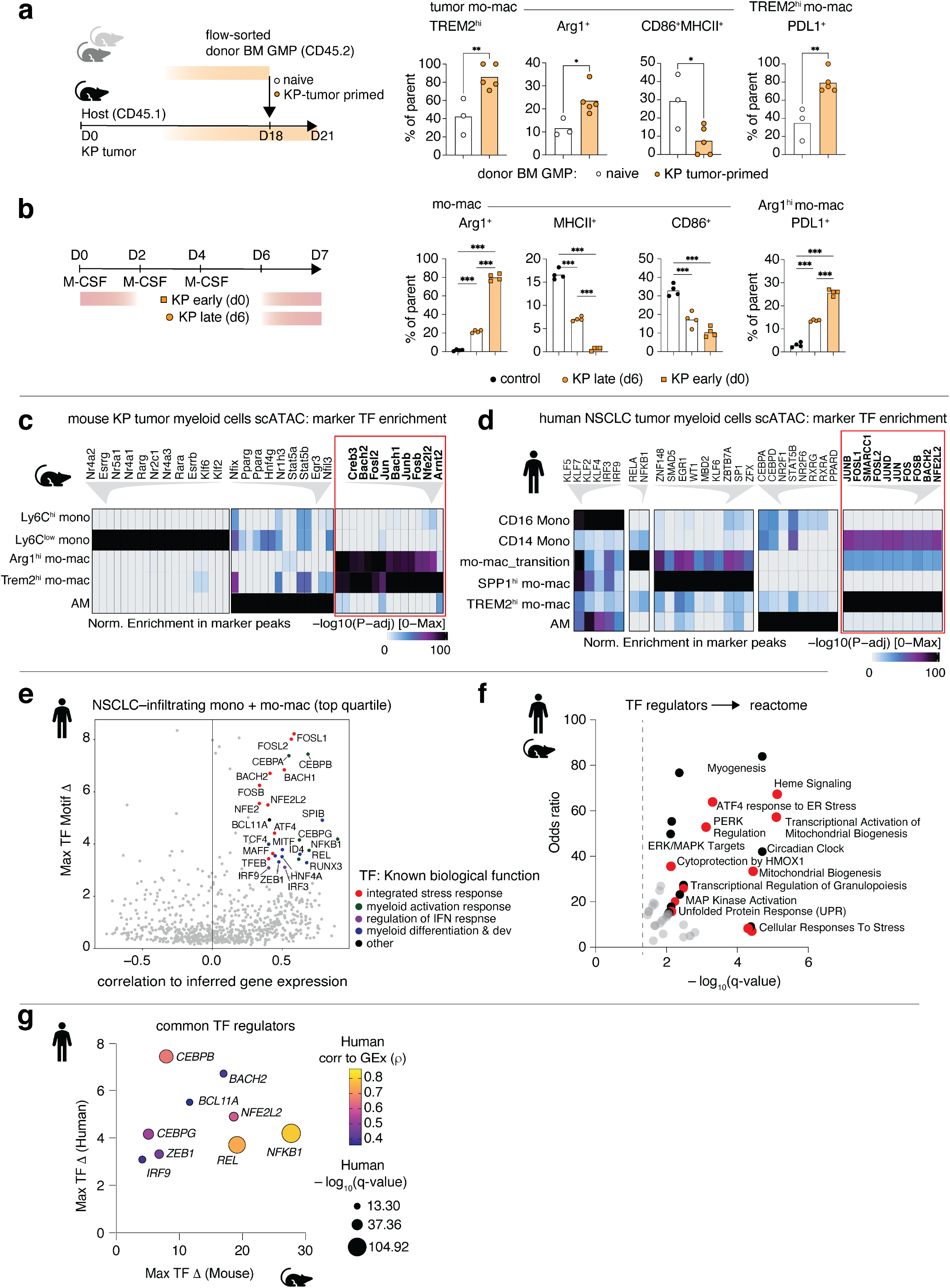
**Tumor myelopoiesis fuels mo-macs in TME with sustained cytoprotective stress responses.** a. *In vivo* tracing of naïve or KP tumor-primed BM GMPs transferred into KP tumor-bearing congenic CD45.1 hosts, with frequency of donor-derived tumor mo-macs expressing GPNMB and CD9 (TREM2^hi^), Arg1 (Arg1^hi^), CD86 and MHCII (CD86^+^MHCII^+^), and frequency of donor-derived TREM2^hi^ mo-macs expressing PDL1. N=3–5 per group. One experiment. b. In vitro BM progenitor-derived macrophage culture with KP tumor conditioning early and late in differentiation or no conditioning (control), with frequency of macrophages expressing Arg1, MHCII, CD86, and frequency of Arg1^hi^ macs expressing PDL1. N=4 per group, representative of two independent experiments. c. scATAC-seq heatmap depicting normalized score for transcription factor (TF) motif accessibility enriched in marker peaks of myeloid cell states in lung of naïve and tumor-bearing mice. N=4 pooled. d. scATAC-seq heatmap depicting normalized score for TF motif accessibility enriched in marker peaks of indicated lung-infiltrating myeloid cell states of patients with NSCLC. N=14 pooled. e. scATAC-seq candidate TF regulators in NSCLC-infiltrating monocyte and macrophage clusters of patients with NSCLC, prioritized by maximum TF motif deviation (ι1) across clusters. Dot color indicating major known biological pathway. f. Reactome pathway terms enriched in TF regulators identified from mouse and human analysis, ranked by adjusted p- value (log q-value) and odds ratio. Red dots linked to stress-associated cytoprotective signaling. g. Prioritization of common top-quartile TF regulators identified from mouse and human analysis, ranked by maximum TF deviation (ι1), and colored by correlation to human TF gene expression (GEx). *p*-values computed by unpaired t-test across conditions (a). *p*-values computed by one-way ANOVA with Sidak’s multiple comparisons test (b). P-value of < 0.05 denoted *; p-values < 0.01 denoted **; p-values < 0.001 denoted ***.

We conducted paired scRNA-seq and scATAC-seq on myeloid-enriched immune cells isolated from naïve and KP tumor-bearing lungs. Building on our previous work^14, 35^, we could identify discrete scRNA-seq clusters of inflammatory Ly6C^hi^ monocytes, patrolling Ly6C^low^ monocytes, resident tissue alveolar macrophages (AM), and major interstitial mo-mac subsets such as Arg1^hi^ mo-macs^34^ and Trem2^hi^ mo- macs^32^ (**Extended Data Fig. 2A**). Using gene-gene correlation^36, 37^, we identified a module of metabolic genes implicated in anti-apoptotic cytoprotection, detoxification, glycolytic shift, and regulation of oxidative stress (e.g.- *Aldoa, Ldha, Pgk1, Sgk1, Pfkp, Ptafr, Got1, Tgm2, Bcl2a1b, Slc7a11, Prdx1, Txnrd1, Hmox1, Vegfa, Socs3*) that was specifically enriched in tumor-infiltrating immunoregulatory Arg1^hi^ mo-macs and Trem2^hi^ mo-macs (**Extended Data Fig. 2B**, gene list in **Supplementary Table 2**). Background-corrected gene set enrichment analyses indicated this co-regulated gene module was regulated by TFs such as *STAT3, NFE2L2, HIF1A, KLF4, SPI1, CEBPB*; reminiscent of our findings in BM progenitors (**Extended Data Fig. 2B**). In the scATAC-seq data, we captured a similar diversity of tumor-infiltrating myeloid cells through independent clustering of chromatin accessibility features (**Extended Data Fig. 2C**) and verified their identity by visualizing known marker gene loci (**Extended Data Fig. 2D**). Upon assessing TF motifs enriched in differentially accessible regions across clusters, we could recapitulate cell-type specific enrichment of lineage-determining TF motifs, such as *Stat5*^38^, *Pparg*^39^ in AMs and *Nr4a1*^40^ in patrolling Ly6C^low^ monocytes (**Fig. 2C**). Importantly, we found that TFs for oxidative stress response such as *Nfe2l2, Fosl2*^41^*, and Bach1*^42^ were specifically enriched in tumor-dominant Arg1^hi^ and Trem2^hi^ mo-macs compared to naïve tissue resident populations such as AMs (**Fig. 2B**). These results, echoing our findings in BM myeloid progenitors, suggest that oxidative stress-induced cytoprotective chromatin changes initiated in tumor-educated myeloid progenitors are maintained along the monocytic lineage in tumoral mo-macs.

### Stress-induced cytoprotective response identified along monocytic lineage in patients with lung cancer

To determine whether the stress-associated chromatin changes observed across the monocytic lineage of tumor-bearing animals was also induced in patients with cancer, we profiled paired single-cell chromatin accessibility and gene expression of all immune cells in human NSCLC primary lung tumors and adjacent lung tissue^43^. The resultant dataset spans 14 patients with NSCLC (metadata in **Supplementary Table 3**) and builds prior work from our group encompassing 35 patients with NSCLC^44^. Based on marker gene-based filtering in our 10x Multiome dataset of 5 patients, we isolated 4,177 myeloid cells and classified them according to their nuclear RNA-seq profiles and by weighted gene correlation network analysis (**Extended Data Fig. 2E** and **2F**; gene list in **Supplementary Table 4**). We could distinguish monocytes and transitional mo-mac states having recently infiltrative monocyte features (mo-mac_transition) from regulatory SPP1^hi^ mo-macs and immunosuppressive TREM2^hi^ mo-macs (**Extended Data Fig. 2F** bottom; gene module in **Supplementary Table 4**). We integrated the cellular identity from Multiome-capture to 24,346 scATACseq–captured myeloid cells from 10 patients and verified the fidelity of gene-activity score markers in these clusters (**Extended Data Fig. 2G**). In close agreement with our mouse analyses, we observed motifs for cytoprotective TFs such as *NFE2L2, FOSL2, JUN, and BACH1* to be enriched in marker peaks of tumor-infiltrating activated CD14 monocytes, mo- mac_transition, and tumoral TREM2^hi^ mo-macs (**Fig. 2D**). Notably, SPP1^hi^ mo-macs had motif enrichment for TGFβ–BMP related TFs such as *EGR1*, *SMAD5*, *ZBTB7A*, and *KLF6*^45, 46^.

Subsequently, we conducted a targeted analysis to identify candidate TF regulators of tumor-associated myelopoiesis with sustained cell type-specific activity in TME monocytes and mo-macs. Encouragingly, putative TF regulators in our mouse dataset included *Spi1/PU.1*, *Mafb,* and *Cebpb* important to mo-mac differentiation and identity^16, 21^, as well as NF-κB/Rel family members *Nfkb1, Relb*, *Hivep3* and AP–1 family members *Jun/Fos* associated with early response to inflammation and growth factors^47^ (**Extended Data Fig. 2H**). Strikingly, we again observed the nexus of TFs known to regulate cytoprotective programs in response to oxidative stress, principally *Nfe2l2, Bach1, Fosl2*, and small MAF members *Mafk, Mafg*. Concurrent analyses in our human NSCLC dataset of TME monocytes and mo-macs indicated very similar candidate regulators such as *NFKB1, REL*, *SPIB*, *CEBPB, CEBPA* but also oxidative stress– and integrated stress–response regulators *NFE2L2, BACH1, MAFF, FOSL2, ATF4* (**Fig. 2E**). Pathways enriched downstream of TF regulators across our mouse and human TME mo-macs included terms surrounding *Heme signaling, Cytoprotection by HMOX1, Response to ER Stress,* and *Mitochondrial Biogenesis* (**Fig. 2F**). Thus, independent analyses across mouse lung cancer and human NSCLC identified a broadly conserved bloc of candidate TFs involved in shaping the fate of monocytic cells infiltrating the lung TME, including TFs we previously observed in tumor-educated BM progenitors. We further prioritized the TFs based on correlation to gene expression in human tissue monocytes and mo- macs, identifying *NFKB1, REL, NFE2L2*, and *CEBPB* (**Fig. 2G**). Given extensive prior work on the role of NF-κB/Rel and C/EBP pathway activation during myeloid cell differentiation in the TME^48–50^, we homed in on cytoprotective TF *NFE2L2* (NRF2).

### Stepwise increase in NRF2 signaling occurs along monocytic lineage in lung cancer

We were especially interested in NRF2 due to its role in driving antioxidant gene batteries, promoting resistance to lipid-associated ferroptosis, and opposing NF-κB pro-inflammatory cascades^51–58^. Our data so-far demonstrated that NRF2 was differentially accessible in tumor-associated BM myeloid progenitors, suggesting priming of stress-responsive pathways prior to mo-mac differentiation in the TME. NRF2 binding has recently been shown to directly influence pro-inflammatory signals by suppressing type-I IFN pathway genes and limit inflammasome activation^59–63^, which also aligns with our observations of IFN hypo-responsiveness in tumor-educated BM myeloid progenitors (**Fig. 1H**). In the lung, we observed relative TF motif deviation for NRF2 to be highest in immunoregulatory Arg1^hi^ and Trem2^hi^ mo-macs that accumulate in late-stage tumors (**Fig. 3A**). We probed the paired single-cell transcriptomic data of tumor-infiltrating myeloid cells and calculated an aggregate score for curated NRF2 downstream genes as a readout of pathway activation^62^. We did not rely on *Nfe2l2* gene expression alone since NRF2 is regulated post-transcriptionally and post-translationally^64^. There was a stepwise increase in pathway activation with differentiation into mo-macs, with Trem2^hi^ mo-macs having the highest NRF2 activation score^65^ (**Fig. 3A**; gene list in **Supplementary Table 1**). Similarly, in our human NSCLC data, *NFE2L2* TF motif deviation was highest in immunosuppressive TREM2^hi^ mo-macs and recently infiltrated mo-mac_transition, with concomitantly high NRF2 activation score indicating downstream gene activation (**Fig. 3B**). As part of corroborating our findings, we calculated the per-patient NRF2 activation score for monocytes and macrophage clusters in an independent scRNA-seq dataset^44^ and found the NRF2 activation score to be highest in tumor-associated TREM2^hi^ mo-macs (**Fig. 3C**). We additionally found the NRF2 activation score to be higher in tumor-infiltrating CD14+ monocytes compared to circulating CD14+ monocytes from the same patients (**Fig. 3D**), suggesting a sustained activation of the gene program upon infiltrating the TME.

**Fig. 3:**
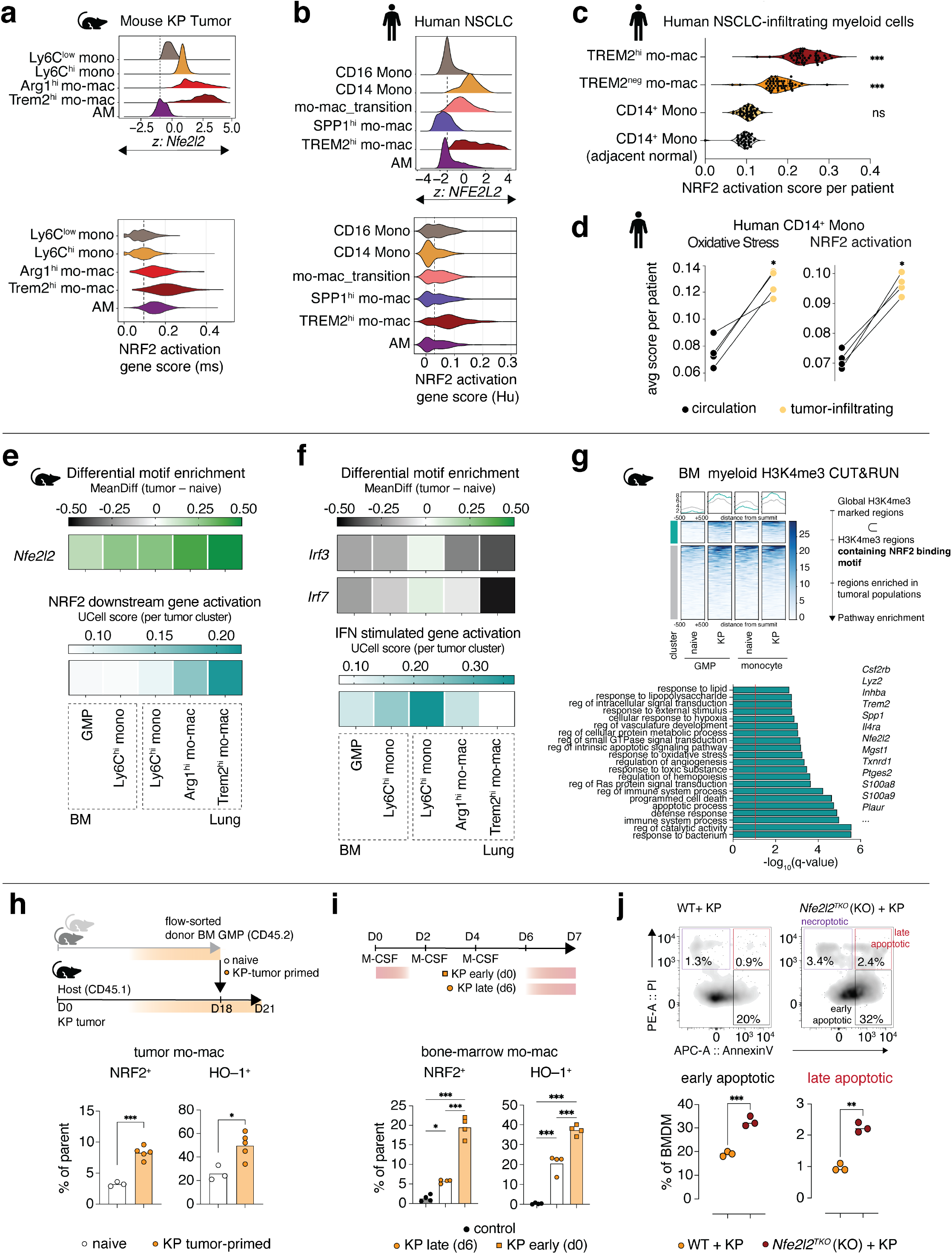
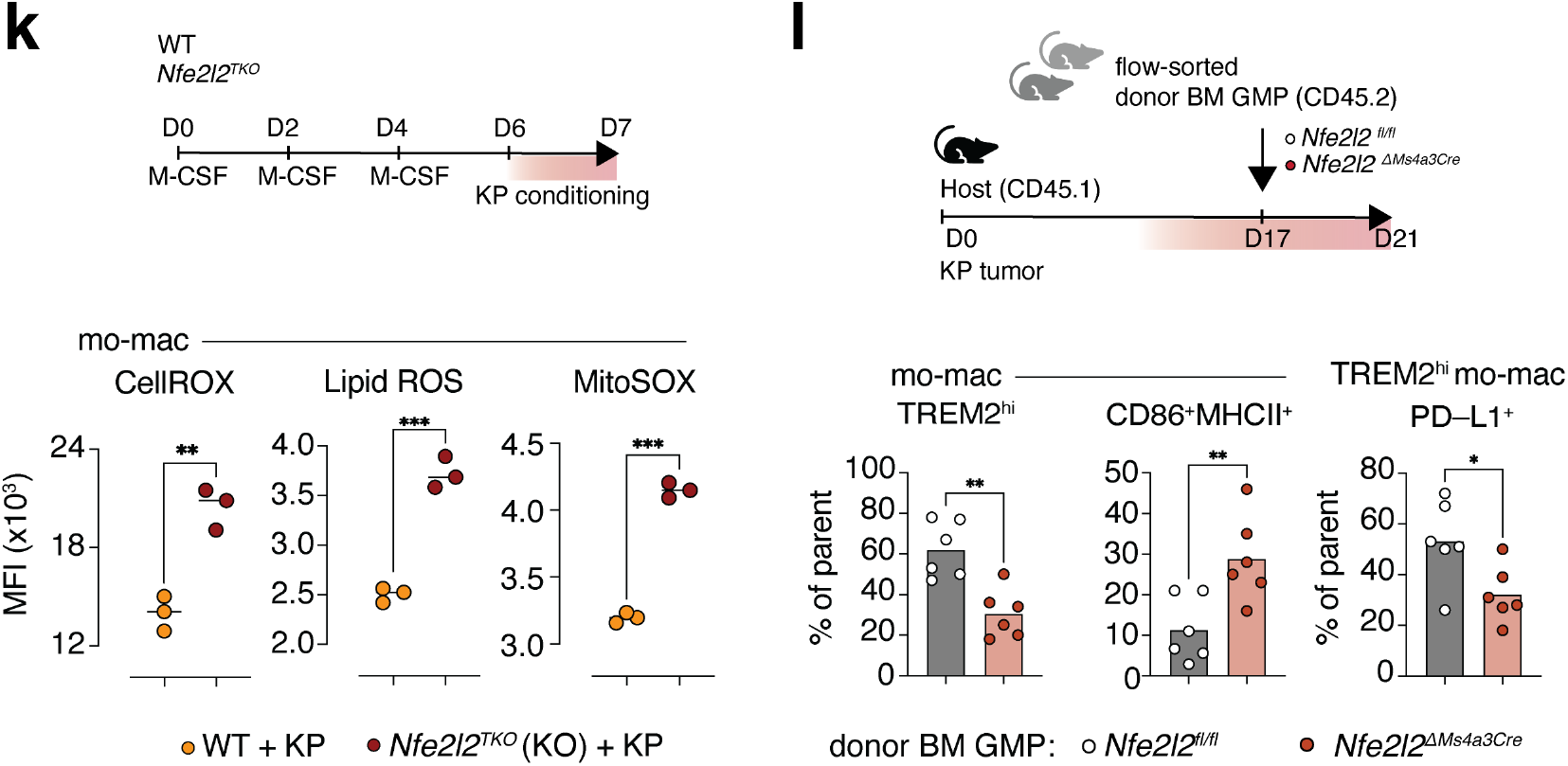
**Stepwise activation of NRF2 signaling regulates tumor-associated mo-mac survival and immunosuppressive function** a. ChromVAR motif deviation for *Nfe2l2* (TOP) and UCell-computed score for NRF2 downstream gene program activation (BOTTOM) in mouse lung tumor-infiltrating myeloid cell clusters. N=4 pooled. b. ChromVAR motif deviation for *NFE2L2* (TOP) and UCell-computed score for NRF2 downstream gene program activation (BOTTOM) in human NSCLC-infiltrating myeloid cell clusters. N=14 pooled. c. NRF2 downstream gene activation score per-patient in human lung-infiltrating myeloid cells from independent validation set of NSCLC re-analyzed from *Leader et al. 2021*. N=35 patients. d. Normalized UCell-computed scores per-patient for Oxidative stress response (LEFT) and NRF2 downstream gene activation (RIGHT) in circulating CD14 monocytes and NSCLC-infiltrating CD14 monocytes for matched patient samples. N=4 patients. e. Relative *Nfe2l2* TF motif enrichment in tumor condition relative to naïve condition (TOP) and UCell-computed NRF2 downstream gene activation in tumor condition (BOTTOM) across indicated myeloid populations. f. Relative TF motif enrichment for *Irf3* and *Irf7* in tumor condition relative to naïve condition (TOP) and UCell-computed IFN stimulated gene program in tumor condition (BOTTOM) across indicated myeloid populations. g. Pathways enriched in indicated cluster of H3K4me3 promoter peaks containing NRF2 binding motif in BM GMPs and Ly6C^hi^ monocytes from KP tumor-bearing mice compared to naïve mice, with exemplar genes indicated. Curated terms arranged by adjusted p-value (log q-value). h. *In vivo* tracing of naïve or KP tumor-primed BM GMPs transferred into KP tumor-bearing congenic CD45.1 hosts, with frequency of donor-derived tumor mo-macs expressing NRF2 and HO-1. N=3–5 per group. One experiment. i. In vitro BM progenitor-derived macrophage culture with KP tumor conditioning early and late in differentiation or no conditioning (control), with frequency of macrophages expressing NRF2 and HO-1. N=4 per group, representative of two independent experiments. j. Representative flow cytometry plots for AnnexinV and Propidium iodide (PI) in NRF2KO (KO) or control (WT) bone- marrow derived macrophages exposed to KP tumor conditioned media, with relative frequency of apoptotic cells. N=3 per group, representative of three independent experiments. k. Relative MFI quantification of CellROX (reactive oxygen species), LiperFluo (lipid peroxidation), and MitoSOX (mitochondrial oxidative burden) in NRF2KO (KO) or control (WT) bone-marrow derived macrophages exposed to KP conditioned media. N=3 per group, representative of three independent experiments. *In vivo* tracing of KP tumor-primed BM GMPs transferred from Nfe2l2^ΔMs4a3^ mice or Nfe2l2^fl/fl^ control littermates into KP tumor-bearing congenic CD45.1 hosts, with frequency of donor-derived tumor mo-macs expressing GPNMB and CD9 (TREM2^hi^), CD86 and MHCII (CD86^+^MHCII^+^), and frequency of donor-derived TREM2^hi^ mo-macs expressing PDL1. N=6 per group. One experiment. *p*-values computed by one-way ANOVA with Dunnett’s multiple comparisons test (c), or Sidak’s multiple comparisons test (i). *p*-values computed by paired t-test (d), or unpaired t-test (h),(j),(k),(l). *p*-values computed by hypergeometric test with multiple test correction (g). P-value of < 0.05 denoted *; p-values < 0.01 denoted **; p-values < 0.001 denoted ***.

Given the persistent NRF2 pathway activation in TME mo-macs with known immunoregulatory function, we next explored when NRF2-linked gene programs are activated during GMP differentiation to monocytes and mo-macs. We analyzed a replicate scRNA- and scATAC-seq integrated dataset filtered on a continuum of monocytic lineages across tumor-associated BM, blood, and lung of late-stage KP tumor–bearing mice. We found *Nfe2l2* TF motif accessibility to again be highest in lung TME Arg1^hi^ and Trem2^hi^ mo-macs (**Extended Data Fig. 3A**). This is consistent with observations that persistent oxidative stress generated in the TME due to accumulation of cellular low-density oxidized lipoprotein can stimulate NRF2 protective pathways^63, 66, 67^. To validate the activation of NRF2-associated programs *in vivo*, we assayed the levels of nuclear NRF2 and intracellular heme oxygenase i.e., HO-1 (encoded by *Hmox1)*^68^ by flow cytometry in myeloid cells infiltrating KP lung tumors. Across two orthotopic models of NSCLC, we found that TREM2^hi^ mo-macs were the main myeloid population in tumors with stable nuclear NRF2 and intracellular HO-1 (**Extended Data Fig. 3B**).

Crucially, we observed tumor-educated BM GMPs and monocytes had increased NRF2 accessibility compared to naïve counterparts, but this was not matched by downstream gene program activation (**Fig. 3E**), implying poising of NRF2-associated gene loci in BM. We also identified a surge of IFN-response genes in tumor-infiltrating monocytes^24^ with subsequent decrease in *Irf3/Irf7* accessibility and lower IFN-stimulated gene program in TME mo-macs (**Fig. 3F**); aligning with work demonstrating negative regulation of Type-I IFN response by NRF2 activation^60, 62^. To further interrogate the poised programs associated with NRF2 in BM progenitors, we assessed H3K4me3 signal in open chromatin regions containing the NRF2 binding motif and differentially increased in tumor-educated BM GMPs and monocytes. Here, we found enrichment of pathways involving *regulation of apoptosis, regulation of hemopoiesis, response to oxidative stress, response to hypoxia,* along with *regulation of immune system process* and *defense response* (genes such as *S100a8/9, Plaur, Inhba, Trem2, Spp1,* and *Il4ra*) (**Fig. 3G**). These data demonstrate TME macrophage-centric gene programs spanning immunoregulation, programmed cell death, and metabolism to be poised in BM progenitors under the control of NRF2, providing unparalleled insight into the mechanics of immunosuppressive ‘pre-conditioning’ in the BM.

### NRF2 is a regulator of mo-mac survival and immunosuppressive function in murine and human tumors

Given our observations of chromatin accessibility for NRF2 binding in BM progenitors with downstream gene activation in tissue mo-macs, we next evaluated whether early sensing of tumor-stress in myeloid progenitors matters for progeny mo-mac NRF2 activation in the TME. To do so, we injected sorted GMPs primed in naïve or KP tumor-bearing mouse BM into congenic hosts bearing late-stage KP tumors. We found that tumor-primed GMPs, after prior exposure to oxidative stress in BM, develop into mo-macs with increased NRF2 activity and HO-1 (**Fig. 3H**). Furthermore, in *ex vivo* BMDM culture models, we observed that early exposure to tumor cues during macrophage differentiation resulted in differentiated BMDMs with increased activation of NRF2 and expression of HO-1 (**Fig. 3I**). These results collectively suggested that initial exposure of BM myeloid progenitors to tumor cues is the ‘first hit’ that primes NRF2-associated oxidative stress pathways and is linked to a more immunoregulatory fate in the TME.

Subsequently, we interrogated the necessity of NRF2 activation for our observed immunoregulatory phenotype in the TME. We cultured BMDMs from NRF2 constitutive knockout mice (referred to as *Nfe2l2^TKO^*) or WT counterparts and exposed them to tumor conditioning to mimic TME polarization. Remarkably, the absence of NRF2 resulted in tumor-educated BMDMs undergoing increased cell death due to increased sensitivity towards lipid peroxidation-linked ferroptosis (**Fig. 3J**). The viable cells from tumor-educated *Nfe2l2^TKO^*BMDMs had reduced expression of HO–1 and immunoregulatory markers Arg1 and PD–L1 with increased expression of costimulatory markers MHCII, CD86, and CD40 (**Extended Data Fig. 3C** and **3D**). Orthogonally, we exposed BMDMs during tumor conditioning to ML385, a validated inhibitor of Nfe2l2-Mafg DNA binding and transcriptional activity^69^ or OB24, a non-competitive inhibitor of HO–1^70^. Addition of ML385 or OB24 in an acute setting resulted in a reduction of HO–1 level as expected, but also caused a phenotypic shift with reduction in Arg1 as well as increased expression of CD86 and MHCII (**Extended Data Fig. 3E**). Importantly, we observed that tumor–educated *Nfe2l2^TKO^* BMDMs had significantly increased ROS burden, lipid peroxidation, and mitochondrial stress (**Fig. 3K**), underlying the increased susceptibility to ferroptosis. These results indicated that NRF2 activation functionally drives cytoprotective resistance to ferroptosis and controls immunosuppressive genes in TME mo-macs.

### NRF2 signaling in myeloid lineage sustains myelopoiesis and promotes immunosuppression in the TME

Considering the important role played by NRF2 in mo-mac survival pathways as well as immunoregulation, we interrogated how loss of NRF2 functionally impacts myelopoiesis and the intratumoral fate of mo-macs. To do so, we generated conditional knockout mice wherein NRF2 is floxed out by Cre recombinase under the *Ms4a3* promoter (hereby referred to as *Nfe2l2^ΔMs4a^*^3^) restricted to granulocytic-monocytic precursors^71^. To ascertain the cell-intrinsic effects of NRF2 loss on myeloid differentiation under acute tumor cues, we adoptively transferred GMPs from CD45.2 *Nfe2l2^ΔMs4a3^*or *Nfe2l2^fl/fl^* control BM into congenic CD45.1 hosts bearing KP tumors. While there was no discernable difference in the number of donor-derived monocytes and mo-macs in the lung TME (**Extended Data Fig. 3F**), we observed GMPs from *Nfe2l2^ΔMs4a3^* mice differentiated into more immunostimulatory CD86^+^MHCII^+^ mo-macs and strikingly fewer TREM2^hi^ mo-macs in the lung TME (**Fig. 3L**), with the TREM2^hi^ mo-macs having decreased PDL1 expression.

Subsequently, we implanted KP lung tumors orthotopically in *Nfe2l2^ΔMs4a3^*or *Nfe2l2^fl/fl^* mice. The lack of myeloid NRF2 resulted in a significant reduction in tumor burden associated with greater overall survival (**Fig. 4A** and **Extended Data Fig. 4A**). Similar results were obtained in the LLC1 aggressive lung cancer model (**Extended Data Fig. 4B)** and the B16-F10 model of lung metastases (**Extended Data Fig. 4C**). Assessing the levels of nuclear NRF2 and expressed HO–1, we could confirm that mo-macs in KP lung tumors of *Nfe2l2^ΔMs4a3^* mice had attenuated NRF2 signaling (**Fig. 4B** and **Extended Data Fig. 4D**). We also generated transgenic mice wherein Kelch-like ECH-associated protein 1 (KEAP1) locus was floxed under Ms4a3Cre resulting in GMP-restricted loss of KEAP1, denoted *Keap1^ΔMs4a3^*. KEAP1 is a component of the Cullin 3 (CUL3)–based E3 ubiquitin ligase complex controlling the stability of NRF2^72^, and deleting KEAP1 in myeloid lineage leads to sustained NRF2 activity. Lung tumors were significantly larger in *Keap1^ΔMs4a3^* mice when compared to negative littermates (**Extended Data Fig. 4E**), supporting our findings in the *Nfe2l2^ΔMs4a3^* experiments. These data collectively suggest that sustained NRF2 signaling in monocytic-lineage cells is crucial for blunting anti-tumor immunity. Reduced tumor burden in *Nfe2l2^ΔMs4a3^* mice was associated with a stark reduction in number of TREM2^hi^ mo-macs in the TME and a compensatory increase in CD86^+^MHCII^+^ antigen presentation proficient mo-macs (**Fig. 4C**). Within TREM2^hi^ mo-macs of *Nfe2l2^ΔMs4a3^* mice, we observed a notably reduced expression of inhibitory PD–L1 (**Fig. 4D**). Reassuringly, the mo-mac phenotype in *Keap1^ΔMs4a3^*mice was consistent with a tumor-promoting role, with more abundant TREM2^hi^ mo-macs in the TME expressing PD–L1 (**Extended Fig. 4F** and **4G**).

**Fig. 4:**
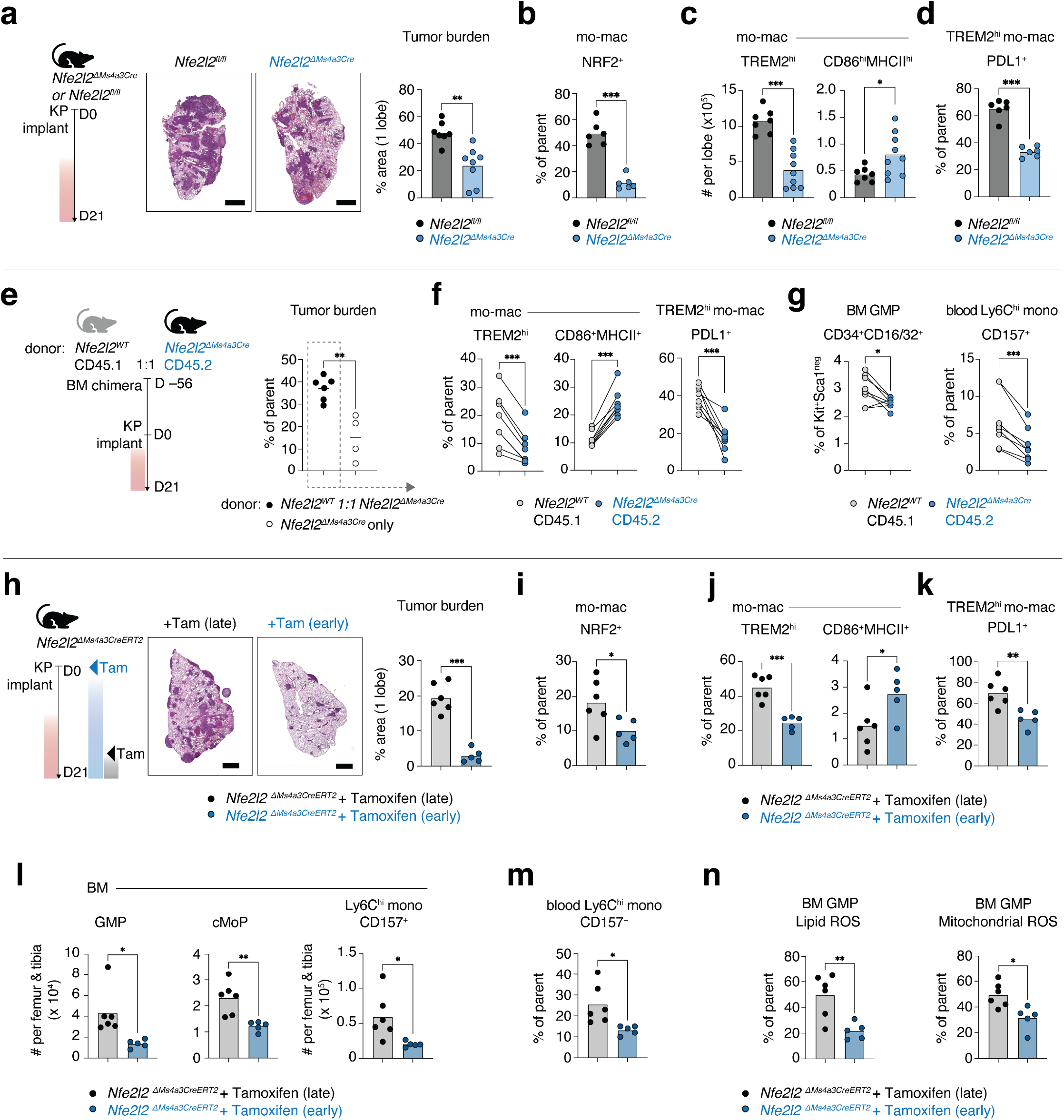
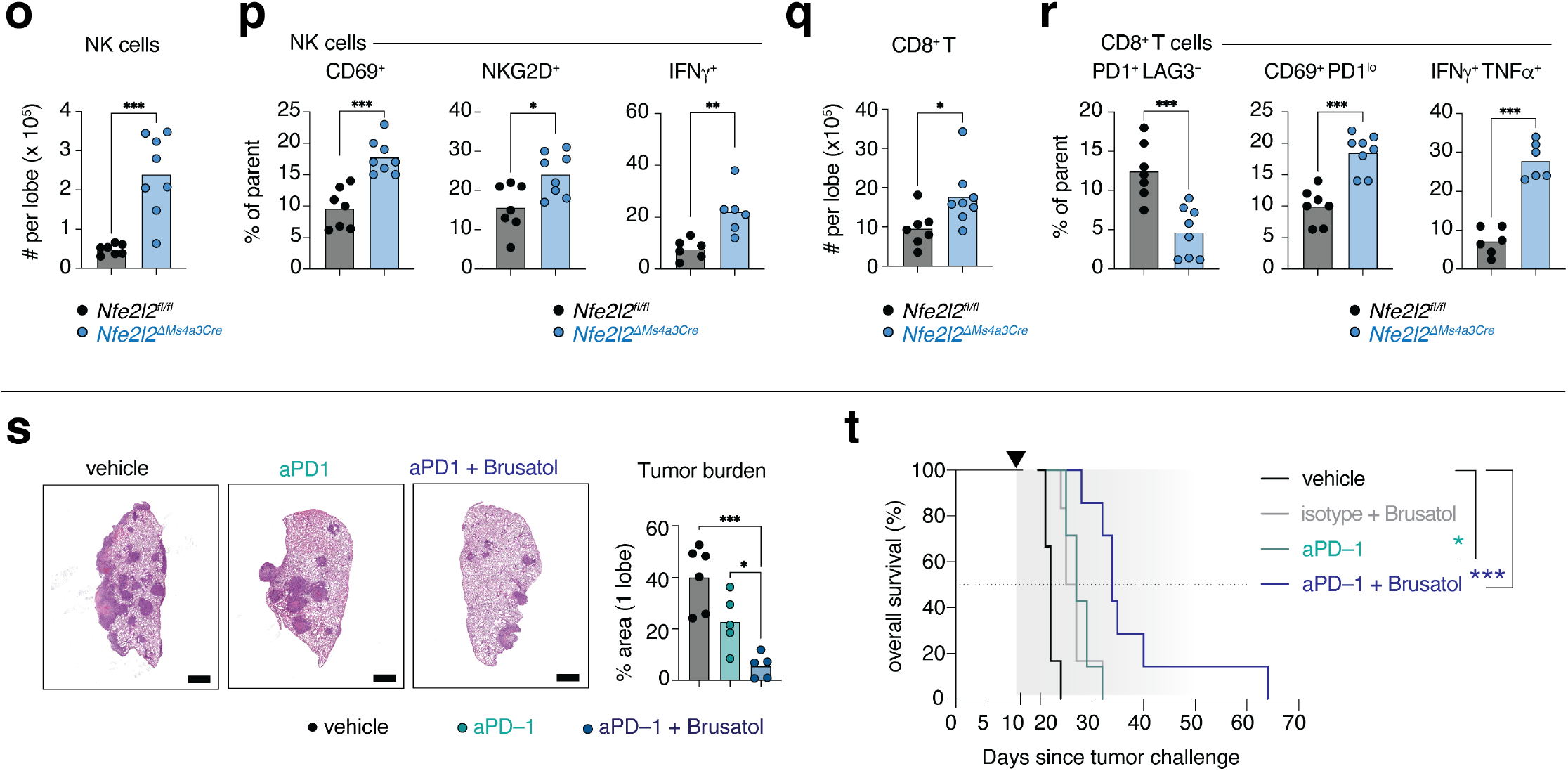
**NRF2 signaling sustains myelopoiesis promoting NK and T cell immunosuppression in TME** a. Representative lungs from KP tumor-bearing Nfe2l2^ΔMs4a3^ mice or Nfe2l2^fl/fl^ negative littermates with quantification of tumor burden. N=7–8 mice per group. Data are individual data points with bar denoting mean. b. Nuclear NRF2 quantified in tumor-infiltrating mo-macs of Nfe2l2^ΔMs4a3^ mice or Nfe2l2^fl/fl^ littermates. N=6 mice per group. c. Number of tumor-infiltrating mo-macs expressing GPNMB and CD9 (TREM2^hi^), CD86 and MHCII (CD86^+^MHCII^+^), in Nfe2l2^ΔMs4a3^ mice or Nfe2l2^fl/fl^ negative littermates. N=7–9 mice per group. d. Frequency of tumor-infiltrating TREM2^hi^ mo-macs expressing immunoregulatory marker PDL1 in Nfe2l2^ΔMs4a3^ mice or Nfe2l2^fl/fl^ negative littermates. N=6 mice per group. e. Schematic of mixed-BM chimera and quantification of tumor burden in Nfe2l2^WT^ 1:1 Nfe2l2^ΔMs4a3^ donor chimera and Nfe2l2^ΔMs4a3^ donor chimera. N=4–6 mice per group. Data are individual data points with bar denoting mean. f. Frequency of lung-infiltrating mo-macs from indicated donors expressing GPNMB and CD9 (TREM2^hi^), CD86 and MHCII (CD86^+^MHCII^+^), and frequency of lung-infiltrating TREM2^hi^ mo-macs from indicated donors expressing immunoregulatory marker PDL1. N=8 mice per group. g. Frequency of BM CD34^+^ myeloid progenitors (LEFT) and blood-circulating CD157^+^ Ly6C^hi^ monocytes (RIGHT) from indicated donors. N=8 mice per group. h. Schematic of Tamoxifen-based early vs late temporal deletion of Nfe2l2 in Nfe2l2^ΔMs4a3CreERT2^ mice, with representative lung tumor burden and quantification of tumor burden. N=5–6 mice per group. Data are individual data points with bar denoting mean. i. Nuclear NRF2 quantified in tumor-infiltrating mo-macs of Nfe2l2^ΔMs4a3CreERT2^ mice with early vs late deletion of NRF2. N=5–6 mice per group. j. Number of tumor-infiltrating mo-macs expressing GPNMB and CD9 (TREM2^hi^), CD86 and MHCII (CD86^+^MHCII^+^), in Nfe2l2^ΔMs4a3CreERT2^ mice with early vs late deletion of NRF2. N=5–6 mice per group. k. Frequency of tumor-infiltrating TREM2^hi^ mo-macs expressing immunoregulatory marker PDL1 in Nfe2l2^ΔMs4a3CreERT2^ mice with early vs late deletion of NRF2. N=5–6 mice per group. l. Abundance of GMPs, cMoPs, and CD157^+^ Ly6C^hi^ monocytes in BM of KP tumor-bearing Nfe2l2^ΔMs4a3CreERT2^ mice with early vs late deletion of NRF2. N=5–6 mice per group. m. Frequency of CD157^+^ Ly6C^hi^ monocytes in circulation of KP tumor-bearing Nfe2l2^ΔMs4a3CreERT2^ mice with early vs late deletion of NRF2. N=5–6 mice per group. n. Relative LiperFluo (lipid peroxidation), and MitoSOX (mitochondrial oxidative burden) in BM GMPs of KP tumor-bearing Nfe2l2^ΔMs4a3CreERT2^ mice with early vs late deletion of NRF2. N=5–6 mice per group. o. Flow cytometry quantification of lung-infiltrating NK cells in KP tumor-bearing Nfe2l2^ΔMs4a3^ mice or negative littermates at day 21. N=7–8 mice per group. p. Frequency of lung-infiltrating NK cells expressing markers CD69, NKG2D, and producing IFNγ in KP tumor-bearing Nfe2l2^ΔMs4a3^ mice or negative littermates. N=7–8 mice per group. q. Flow cytometry quantification of lung-infiltrating CD8^+^ T cells in KP tumor-bearing Nfe2l2^ΔMs4a3^ mice negative littermates at day 21. N=7–8 mice per group. r. Frequency of lung-infiltrating CD8^+^ T cells expressing inhibitory markers PD1, LAG3, activating markers CD69, and producing IFNγ and TNFα in tumor-bearing Nfe2l2^ΔMs4a3^ mice or littermates. N=7–8 mice per group. s. Representative histology of KP tumor-bearing mice treated with Brusatol (NRF2 inhibitor) in conjunction with anti-PD1 immunotherapy, anti-PD1 alone, or vehicle; with quantification of tumor burden. N=5–7 mice per group, representative of two independent experiments. t. Kaplan-Meier plot depicting overall survival of KP tumor-bearing mice treated with Brusatol (NRF2 inhibitor) in conjunction with anti-PD1 immunotherapy, Brusatol alone, anti-PD1 alone, or vehicle. N=6–8 mice per group. One experiment. *p*-values computed by unpaired t-test (a)–(e),(h)–(r), and paired t-test (f),(g). *p*-values computed by one-way ANOVA with Dunnett’s multiple comparisons test (s). p-values computed by Log-rank (Mantel-Cox) test (t). P-value of < 0.05 denoted *; p- values < 0.01 denoted **; p-values < 0.001 denoted ***.

While these observations underscore the clear impact of NRF2 signaling in driving mo-mac fate in the TME, they do not reveal the full extent to which NRF2 poising impacts myelopoiesis. To disentangle local reprogramming in the TME from replenishment due to pathogenic myelopoiesis, we created mixed-BM chimeras with 1:1 reconstitution of CD45.2 *Nfe2l2^ΔMs4a3^* and age-matched NRF2-proficient CD45.1 *Nfe2l2^WT^* mice. This system enabled us to implant KP tumors and study the cell-intrinsic impact of NRF2 loss independent of tumor burden differences. The tumor burden in chimera mice was higher than control mice which received only CD45.2 *Nfe2l2^ΔMs4a3^* BM (**Fig. 4E**). Tumoral TREM2^hi^ mo-macs of *Nfe2l2^ΔMs4a3^*origin were less abundant in the chimera TME and had lower immunosuppressive PD-L1 while CD86^+^MHCII^+^ mo-macs were increased (**Fig. 4F**). Importantly, there was decreased mobilization of *Nfe2l2^ΔMs4a3^* mouse–derived CD34^+^CD16/32^+^ myeloid progenitors in BM and CD157^+^ GMP-origin Ly6C^hi^ monocytes in peripheral blood (**Fig. 4G**), suggesting a tumor burden-independent impact of NRF2 signaling on myelopoiesis at the medullary site.

Additionally, we determined the temporal importance of ablating NRF2 signaling in myeloid precursors prior to or after tumor exposure– by utilizing the tamoxifen-inducible Ms4a3CreERT2 strategy^71^ to generate *Nfe2l2^ΔMs4a3CreERT2^* mice. Tamoxifen administration at the time of KP tumor implantation (i.e. early) in *Nfe2l2^ΔMs4a3CreERT2^* mice resulted in a significant reduction in tumor burden when compared to tamoxifen administration at later stages of progression (**Fig. 4H**). In *Nfe2l2^ΔMs4a3CreERT2^* mice with NRF2 signaling attenuated early, TME mo-macs had lower activation of nuclear NRF2 (**Fig. 4I**), were more immunostimulatory (**Fig. 4J**), and had reduced PD–L1 expression (**Fig. 4K**). Early attenuation of NRF2 signaling in *Nfe2l2^ΔMs4a3CreERT2^* mice i.e., prior to tumor-induced myelopoiesis, was associated with a decrease in BM GMPs, cMoPs, and Ly6C^hi^ monocytes (**Fig. 4L**) with reduced mobilization of CD157^+^ Ly6C^hi^ monocytes into circulation (**Fig. 4M)**. In line with our hypotheses, we found tumor-primed BM GMPs with continuous attenuation of NRF2 signaling had lower lipid peroxidation and mitochondrial oxidative stress (**Fig. 4N**).

Functionally, changes in the myeloid compartment upon myeloid knockout of NRF2 signaling were associated with significant infiltration of NK cells into lung tumors (**Fig. 4O**). NK cells in the lung tumors of *Nfe2l2^ΔMs4a3^* mice had increased CD69 and NKG2D with increased IFNγ secretory capacity, indicating activated tumoricidal qualities (**Fig. 4P**). CD8^+^ T cells were also more abundant in the tumors of *Nfe2l2^ΔMs4a3^*mice (**Fig. 4Q**), exhibiting an activated effector phenotype characterized by reduced exhaustion marker LAG3, increased CD69, and production of IFNγ and TNFα (**Fig. 4R**). Antibody-based depletion of NK cells in tumor–bearing *Nfe2l2^ΔMs4a3^* mice resulted in an increased tumor burden, highlighting the mode of mo-mac immunosuppression to be dependent on NK-cell exclusion (**Extended Data Fig. 4H**). In conclusion, the loss of NRF2 in monocytic-lineage cells resulted in profound changes in the TME, reducing the influx of mo-macs with immunosuppressive phenotype and shifting the balance towards functionally anti-tumor effector cells. Overall, these data suggest that sustained monocytic survival and mo-mac residency in the lung TME driven by activation of NRF2 downstream signaling curtails NK cell-driven tumor surveillance and control^33, 73, 74^.

### Targeting myeloid NRF2 pathway shifts intratumoral macrophage distribution and enhances immunotherapy responses

Given the impact of NRF2 activation on mo-mac immunosuppressive fate curtailing anti-tumor NK cell immunity, we were interested to study if and how myeloid-intrinsic NRF2 pathways influences immunotherapy response. Spurred by our on *in vivo* results around genetic knockout of *Nfe2l2* in myeloid cells, we tested pharmacological inhibition of the NRF2 pathway in a treatment setting. After challenging mice with KP tumors, we administered the quassinoid agent Brusatol^75^ with or without anti-PD1 checkpoint therapy starting at day 9 post-implant. Tissue profiling at 12 days post-treatment suggested robust anti-tumor response and control of tumor burden associated with combination targeting of NRF2 signaling and checkpoint blockade (**Fig. 4S**). We observed that Brusatol and anti-PD1 combination treatment was safe and had a distinct survival advantage beyond the benefit derived from anti-PD1 monotherapy (**Fig. 4T**). We observed a clear decrease in the abundance of mo-macs (**Extended Data Fig. 4I**) along with a significant decrease in frequency of PDL1^+^ TREM2^hi^ mo-macs and concomitant increase in CD86^+^MHCII^+^ mo-macs (**Extended Data Fig. 4J**). There was a substantial influx of NK cells upon combination treatment, with increased expression of CD69 and tumoricidal IFNγ (**Extended Data Fig. 4K** and **4L**). Simultaneously, there was increased infiltration of CD8^+^ T in combination-treated tumors, with the CD8^+^ T cells lacking exhaustion marks PD1 and LAG3, while producing IFNγ and TNFα (**Extended Data Fig. 4M** and **4N**). Overall, we demonstrate the substantial impact of targeting NRF2 signaling on tumor-induced myelopoiesis and mo-mac fate in the TME. Such a myeloid-directed therapeutic strategy results in an unleashing of NK cell and CD8^+^ T cell activity to drive effective anti-tumor immune responses in the lung.

## DISCUSSION

Given that tumor-associated mo-macs associated with immunosuppression & poor prognosis are mostly derived from BM myeloid progenitors, we posited that (1) enhanced myelopoiesis in tumor-bearing hosts influences mo-mac dysfunction, and (2) targeting upstream processes that promote pathogenic myelopoiesis and mo-mac persistence is more effective than merely targeting differentiated mo-macs in the TME. Thus far, the exact contribution of epigenetic changes in BM HSCs and myeloid progenitors to TME mo-mac programs and behavior have never been fully addressed. Leveraging chromatin and transcriptional mapping of tumor-educated myeloid progenitors, monocytes, and tumor infiltrating mo- macs, our work identifies a pivotal point of epigenetic alteration that is initiated in myeloid progenitors and increases along myeloid lineage to promote mo-mac survival and dampen pro-inflammatory pathways in the TME. Our research underscores the importance of identifying and targeting molecular ‘hits’ associated with tumor myelopoiesis in myeloid progenitors. NRF2 activity poising is a consequential important first hit in the BM that is further solidified within the TME (**Extended Data Fig. 4O**).

Our findings in BM myeloid progenitors are indicative of hormetic oxidative stress during steady-state hematopoiesis becoming dysfunctional during chronic inflammation and malignancy^76, 77^. We hypothesize that in tumor-bearing hosts, emergency myeloid-biased expansion at the expense of self-maintenance results in a maladaptive progenitor state wherein progeny cells withstand increased stress on mitochondrial and ER protein machinery via activation of NRF2 and associated pathways^78^. We find NRF2 downstream gene programs to become progressively activated in tumor-infiltrating mo-macs, providing these cells with pro-survival cues and detoxification machinery in the lipid-laden and hypoxic TME. Moreover, our observations of cytoprotective pathway priming could help explain how mo-macs persist during chemo-radiation therapies that elicit tumor cell death and peroxidative stress^79, 80^, and how ‘reactive’ myelopoiesis could contribute to treatment resistance and disease recurrence^81^. Our findings align with work that demonstrates similar stress-induced metabolic adaptations in tumor-associated neutrophils via the enzyme Acod1^82^, as well as mechanisms described for deterministic reprogramming of neutrophils in hypoxia-stressed TME to adopt pro-angiogenic function^83^. It is thus likely that oxidative stress-based cytoprotection against ferroptosis is a conserved mechanism that maintains TME-infiltrative GMP-derived monocytes and neutrophils. Our study illuminates a unified mechanism by which multiple myeloid cell states, supported by untimely mobilization of immature myeloid progenitors, engender immunosuppressive activity in the TME^84, 85^. Importantly, our data extends this paradigm, positing that stress-induced maladaptation of NRF2 signaling is initiated early in tumor-educated BM myeloid progenitors and solidified further along the monocyte and mo-mac lineages in the TME. Future studies will help establish the cell-intrinsic and -extrinsic metabolic implications of such pro-survival pathways in persistent TME mo-macs. Our work also insinuates establishment of epigenetic memory in HSCs^86, 87^ and myeloid progenitors driven by tumor inflammation. Assessing the heritability of chromatin changes wrought by distal tumor cues in mobilized progenitors can provide us with important biomarkers in blood PBMCs for tumor detection, monitoring, and patient stratification.

Notably, our data highlights the feasibility of targeting oxidative stress regulators such as NRF2 in influencing monocyte fate and restoring mo-mac immunogenicity. Targeting NRF2 signaling in myeloid progenitors and TME myeloid cells can complement cytotoxic therapies targeting the NRF2/ KEAP1 axis vulnerability in lung cancers. Indeed, NRF2 hyperactivation is the third most frequent genomic event in LUAD, primarily occurring via KEAP1 and ubiquitin ligase CUL3 loss-of-function mutation^64, 88^. Targeting upstream regulators of cytoprotective and anti-inflammatory genes, such as in our study, can arguably have more wide-ranging and durable impact than targeting genes such as HO-1^68^, by altering the trajectory of monocytic differentiation in the TME to facilitate differentiation into immunostimulatory mo- macs. Therapeutic modalities including TF–targeted protein degrader PROTACs^89^ can accelerate such

endeavors. It is important to note that NRF2 is part of a constellation of cytoprotective stress response regulators including FOSL2, BACH1, ATF4, NFIL3, and DDIT3/CHOP^56, 90–92^, which were also identified in our analyses and likely impinge on metabolic adaptations via PKR-like ER kinase (PERK) activation^93, 94^. Future studies can help decipher how these TFs interplay with each other and NRF2 in poising cytoprotective and immunoregulatory gene loci in myeloid progenitors to orchestrate distinct TME mo-mac phenotypes.

## ACKNOWLEDGEMENTS

M.M. was supported by National Institutes of Health (NIH) grants CA257195, CA254104 and CA154947. S.H. was supported by the National Cancer Institute (NCI) fellowship K00CA223043. B.Y.S. was supported by NIH Medical Scientist Training grant T32GM146636. R.M. was supported by the AACR-AstraZeneca Immuno-oncology fellowship 21-40-12-MATT. We thank the patients and their families for participating in the clinical study. This work was supported by grant R24-072073 to the Immunological Genome Project (ImmGen) consortium. We thank members of the Merad laboratory and Brown laboratory at the Marc and Jennifer Lipschultz Precision Immunology Institute at Mount Sinai for insightful discussions and feedback. We would like to thank Emily Bernstein, Dan Hasson, and Dan Filipescu for their technical advice and feedback. We acknowledge the Human Immune Monitoring Center, the Mount Sinai Biorepository and Pathology Core, and the Mount Sinai Cytometry Core for extensive support and resources. This work was supported in part through the computational resources and staff expertise provided by the Bioinformatics for Next Generation Sequencing (BiNGS) Shared Resource Facility funded by NCI P30 Cancer Center and NIH/NIAMS P30 Skin Biology Disease Resource-based Center support grants. This work was also supported in part through the computational and data resources and staff expertise provided by Scientific Computing and Data at the Icahn School of Medicine at Mount Sinai and supported by the Clinical and Translational Science Awards (CTSA) grant UL1TR004419, and the Office of Research Infrastructure award S10OD026880 and S10OD030463. The content is solely the responsibility of the authors and does not necessarily represent the official views of the NIH.

## AUTHOR CONTRIBUTIONS

Conceptualization: S.H. and M.M. Methodology: S.H., B.G., A.Magen, S.M., A.M.T., and M.M.; Investigation: S.H., J.L., R.M., M.D.P., A.Marks, M.Belabed, P.H., T.C., L.T., K.A., T.D., and G.C. Computational investigation: S.H., B.G., B.Y.S., L.H., A.Magen, B.K., D.D., M.D.P., J.L.L., D.A., and M.Bale Writing – Original Draft: S.H. Writing – Reviewing & Editing: S.H., B.Y.S., and M.M. Resources: S.K–S., R.F., A.J.K., F.G., S.Z.J., S.M., A.M.T., T.U.M., B.D.B., and M.M. Funding and Supervision: M.M.

## SUPPLEMENTARY TABLES

Supplementary Table 1: Mouse and Human gene signatures Supplementary Table 2: Mouse gene module list – BM and Lung Supplementary Table 3: Human cohort metadata Supplementary Table 4: Human gene module list – Lung

Supplementary Table 5: Antibodies list for Mouse and Human flow cytometry Supplementary Table 6: Accession code for public datasets

## MATERIALS & CORRESPONDENCE

Further information and requests for reagents should be directed to the corresponding author, Miriam Merad

## METHODS

### Mice

C57BL/6 mice were obtained from Charles River Laboratories (Wilmington, MA) or Jackson Laboratory (Bar Harbor, ME). Ms4a3^Cre^ mice were a gift from Florent Ginhoux, and subsequently purchased from Jackson laboratory (C57BL/6J-Ms4a3em2(cre)Fgnx/J RRID:IMSR_JAX:036382), and Ms4a3^CreERT2^ mice were received from Florent Ginhoux. Nfe2l2^fl/fl^ floxed mice were purchased from Jackson laboratory (C57BL/6-Nfe2l2tm1.1Sred/SbisJ RRID:IMSR_JAX:025433). NRF2 constitutive KO mice were purchased from Jackson laboratory (B6.129X1-Nfe2l2tm1Ywk/J RRID:IMSR_JAX:017009). Keap1^fl/fl^ floxed mice were purchased from Jackson laboratory (B6(Cg)-Keap1tm1.1Sbis/J RRID:IMSR_JAX:037075). Tumor implantations and other experiments were conducted in mice between 10–14 weeks of age. Both male and female mice were used, and we observed no significant differences between sexes in any experiment. Where applicable, littermate controls were used to minimize variation between mouse strains. Mice were housed in individually ventilated cages at the Mount Sinai specific-pathogen-free (SPF) facilities, provided food and water *ad libitum*, with conditions maintained at 21∼23C and 39∼50% humidity and 12/12 hour dark/light cycle. All experiments were approved by, and in compliance with the Institutional Animal Care and Use Committee of the Icahn School of Medicine at Mount Sinai.

### Human Subjects

Informed consent was obtained utilizing the Universal Consent for Mount Sinai Biorepository (Human Subjects Electronic Research Applications 20-01197), in accordance with the protocol reviewed and approved by the Institutional Review Board (IRB) at the Icahn School of Medicine at Mount Sinai (ISMMS). Participants provided written consent to analysis of their blood and resected tissue. Samples of tumor and non-involved lung were then obtained from surgical specimens of the participants undergoing resection at the Mount Sinai Hospital (New York, NY) in collaboration with the Thoracic Surgery Department, the Mount Sinai Biorepository and Department of Pathology. Analysis of the tumor and non-involved lung samples were performed under IRB Human Subjects Electronic Research Applications 10-00472A, in accordance with the protocol reviewed and approved by the IRB at ISMMS.

### Murine Tumor models

Unlabeled or GFP-transduced *Kras*^LSL-G12D/+^; *Trp53*^fl/fl^ (KP) and non-fluorescent *Kras*^LSL-G12D/+^*; Trp53*^fl/fl^; *Rosa26^A3Bi^*; *Rag1*^-/-^ (KPAR) cells derived from previously reported and validated mouse models of NSCLC ^9, 10^ were used for tumor implantation models. KP lines were maintained at 37C in RPMI supplemented with 10% v/v Fetal Bovine Serum (FBS) and 1% v/v Penicillin/Streptomycin (Pen/Strep), and KPAR lines were maintained at 37C in DMEM supplemented with 10% Fetal Bovine Serum and 1% Pen/Strep. LLC1 carcinoma cells were a gift from Dr. Lucas Ferrari de Andrade at Mount Sinai. B16-F10 melanoma cells were purchased from ATCC (CRL-6475) and maintained in 37C in DMEM with 10% Fetal Bovine Serum and 1% Pen/Strep. All cell lines were screened every 6 months for mycoplasma contamination. Cells were injected *in vivo* when in log-phase of growth and within 3-4 passages of thawing. Depending on the experiment, 500,000 KP cells, 150,000 KPAR cells, 500,000 B16-F10, or 500,000 LLC1 cells were injected intravenously (i.v.) through the tail vein. For survival studies, mice were sacrificed when they exhibited >15% body weight loss or moribund status (labored breathing, hunched posture, cachexia) according to predetermined humane endpoints. For profiling studies, mice were sacrificed at the timepoints described in text. All experiments were approved and in compliance with the Institutional Animal Care and Use Committee of the Icahn School of Medicine at Mount Sinai.

### In vivo treatments

To deplete NK cells specifically *in vivo*, tumor-bearing mice were administered anti-NK1.1 depleting antibody (BioXcell Clone PK136, Cat #BE0036) or appropriate IgG2a isotype control (BioXcell, Cat #BE0085) at indicated time point and continued every other day. Similarly, mice were administered anti- CD8a depleting antibody (BioXcell Clone 2.43, Cat #BE0061) or appropriate IgG2b isotype control (BioXcell, Cat #BE0090) every other day. To assess pharmacological inhibition of NRF2 pathway *in vivo*, tumor-bearing mice were given Brusatol (MedChemExpress Cat #HY-19543) orally, dissolved in 5% DMSO and 95% corn oil. Mice were administered 100 ug of anti-PD-1 neutralizing antibody (BioXcell Clone RMP1-14, Cat #BE0146) i.v. with or without 50 ug of Brusatol every other day starting at day 10 after tumor implantation.

### Tissue processing (Mouse)

Mice were euthanized by CO2 inhalation and death confirmed by cervical dislocation. Mice were subjected to transcardiac perfusion with cold PBS and relevant tissue extracted for downstream studies. Mouse lung lobes were digested on a shaker in RPMI media containing 10% FBS, Collagenase IV (Sigma) and DNase I (Sigma) for 30 minutes at 37C before being triturated through an 18G needle and filtered through a 70 μm mesh. Samples were subjected to RBC Lysis Buffer (BioLegend) for 2 mins at RT and quenched with ice-cold FACS Buffer (phosphate-buffered saline supplemented with 1% bovine serum albumin and 2mM EDTA) prior to downstream processes. Bone marrow was flushed with cold FACS Buffer using a 27G needle from both long bones (femur and tibia) and filtered through a 70 μm mesh. Samples were subjected to RBC Lysis Buffer (BioLegend) for 2 mins at RT and quenched with ice-cold FACS Buffer prior to downstream processes. For assays such as low-input RNA and ATAC seq, marrow cells were enriched for Kit^+^ progenitors using the Mojosort Mouse Lin-neg enrichment kit (BioLegend #480004). Blood was collected by cardiac puncture in EDTA-coated tubes and RBCs were lysed in two successive cycles of 5 mins each, at RT in RBC Lysis Buffer (BioLegend). Samples were quenched with FACS Buffer and kept cold for downstream processes. Where relevant, serum was collected by coagulating blood in regular microcentrifuge tubes for 30 mins at RT prior to centrifugation at 5000g for 15 mins at RT. Serum aliquots were made and stored at -80C until experiment.

### Histology

Tumor-bearing lungs were analyzed at indicated timepoints as follows: the left lung lobe was fixed in 4% paraformaldehyde at 4C, embedded in paraffin, and examined as 5 µm cross-sections. Following hematoxylin and eosin (H&E) staining, lung tissue sections were scanned on slides at 20X magnification using a Leica Aperio AT2 digital scanner and quantified by manual annotation of blinded slides using the Panoramic viewer and QuPath software v0.4^95^.

### Ex vivo culture models

Bone marrow–derived macrophages (BMDMs) were generated ex vivo using established protocols ^33^. Briefly, bone marrow was flushed using cold sterile PBS in sterile conditions (under laminar flow) and RBC-lysed for 1 min at RT. Cells were plated in DMEM containing 10% v/v FBS and 10 ng/ml recombinant M-CSF (Peprotech #315-02). Cells were plated at a concentration of ∼150,000 cells per cm2 on non-treated Petri plates. At day 2, media was replenished 1:1 with fresh media containing 10 ng/ml M-CSF. At day 4, media was replaced with fresh media containing 10 ng/ml M-CSF. At day 6, cells were gently replated onto test plates using ice-cold PBS containing 5 mM EDTA. KP conditioned media (CM) was added at 1:1 ratio with existing media at indicated timepoint. Based on the experimental timepoint, at day 7 or 8, BMDMs (verified 90∼95% of culture condition based on F4/80 and CD11b expression) were gently lifted off plates using ice-cold PBS containing 10 mM EDTA and subjected to downstream processes (flow cytometry, sequencing). KP CM was obtained from sub–confluent tumor cells grown in DMEM containing 10% v/v FBS, spun down to remove large cellular debris and frozen in aliquots at -20C until usage. For experiments involving inhibitors of specific pathways, day 7 BMDMs were exposed to ML385 (MedChemExpress, Cat#HY-100523) or OB24 (MedChemExpress, Cat#HY-118487) at indicated concentration and incubated for 18-24 hours prior to wash off.

### Methylcellulose and Liquid culture assays

Total hematopoietic cells were extracted from indicated mouse BM by flushing one femur with PBS, red blood cell lysed, and cells resuspended to a concentration of 300,000 cells per ml in IMDM (Cytivia) containing 1% pen/strep and 2% FBS. A volume of 0.4 ml of the resultant cells was added to pre-aliquoted 4 ml MethoCult tubes (StemCell Cat#M3434 containing recombinant mouse (rm)SCF, rmIL3, rmIL6, recombinant human (rh)EPO, rhInsulin and Transferrin). The mixture was vortexed and dispensed onto 35-mm culture dishes in triplicates following manufacturer’s instructions. The dishes were incubated in a humidified incubator at 37C, 5% CO2. Colonies were manually counted on day 8 on a gridded scoring dish and averaged across 4 independent plates. Liquid cultures from indicated mouse BM were generated as follows; 500 myeloid progenitors were sorted in triplicates into a 96–well non–TC coated plate (Greiner) containing 150uL IMDM media containing 5% FBS, 1% pen/strep, 50 μM beta-mercaptoethanol + cytokines (25 ng/ml each of rmSCF, IL11, FLT3, and rhTPO and 10 ng/ml each of IL3, GMCSF, and rhEPO). 160 ul of fresh media was replenished every 2 days, and cells were counted using a Countess on day 4, day 6, and day 8 post sort.

### Tissue processing (Human)

Human NSCLC lung tissues were rinsed in cold PBS, minced, and incubated on a shaker for 35 minutes at 37C in RPMI media containing Collagenase IV at 0.25 mg/ml, Collagenase D at 200 U/ml and DNAse– I at 0.1 mg/ml (Sigma). Cell suspensions were then quenched in ice-cold FACS Buffer, triturated through a 18G needle, and filtered through a 70 μm mesh prior to RBC lysis for 2 mins at RT. Cell suspensions were enriched for CD45^+^ cells by either bead selection (bound fraction from Stem Cell EasySep Human CD45 Depletion Kit II) per kit instructions or FACS sorting on a BD FACSAria or Beckman CytoFlex SRT sorter prior to processing for scRNA-seq, scATAC-seq or Multiome.

Human NSCLC blood was processed as follows– PBMCs were isolated by Ficoll gradient and underwent RBC lysis for 5 mins at RT. Cell suspensions were enriched for all myeloid cells or CD34^+^ myeloid cells by either bead selection (bound fraction from StemCell Technologies Custom negative selection Kit or CD34 positive selection II Kit) or FACS sorting on a BD FACSAria or Beckman CytoFlex SRT sorter prior to processing for scRNA-seq, scATAC-seq or Multiome.

### Flow cytometry and fluorescence-activated cell sorting (FACS)

<u>Mouse Tissue</u>: Single cell suspensions from mouse Lung, Blood, and BM were resuspended at desired cellular concentration in ice-cold FACS buffer and subjected to immunostaining in the following ways– cells were first incubated with Fixable Blue Live/Dead Dye (Thermo Fisher Scientific) and CD16/CD32 (clone 93, Biolegend) for 15 mins on ice prior to surface staining. Cells were stained for surface markers for 25 mins on ice (antibody details listed in **Supplementary Table 5**). Subsequently, cells were acquired fresh on BD LSR Fortessa analyzer. Alternatively, cells were fixed using BD Cytofix kit (BD #554655) following manufacturer’s instructions and acquired on analyzer within 3 days of fixation. For experiments assaying intracellular markers, cells were fixed using BD Cytofix/Cytoperm (BD Cat#554722) following manufacturer’s instructions and stained for intracellular antigens in Perm Buffer for 30 mins at 4C. For cytokine staining, cells were first simulated in 10 μg/ml Brefeldin A, 0.2 μg/ml Ionomycin and 0.5 μg/ml PMA (Thermo Fisher Scientific) for 4 hours at 37C prior to intracellular staining. For experiments assaying intranuclear transcription factors, cells were fixed and permeabilized using Ebioscience FOXP3 kit (Thermo Fisher Scientific #00-5523-00) following manufacturer’s instructions and stained for intracellular antigens in Perm Buffer for 30 mins at 4C. (antibody details listed in **Supplementary Table 5**). Subsequently, cells were acquired on BD LSR Fortessa analyzer within 3 days of fixation. For sequencing purposes, cells were stained as above and sorted on BD FACS Aria sorter or Beckman CytoFlex SRT sorter using DAPI or 7-AAD to exclude dead cells. Cells were sorted into pre-chilled FBS-coated microcentrifuge tubes in 200 μl of PBS containing 0.5% BSA. To assay reactive oxygen species and lipid peroxidation stress, cells were stained with CellROX (Invitrogen Cat#C10444) and LiperFluo (Dojindo, Cat#L248) according to manufacturer’s instructions. To assess mitochondrial function and membrane polarity, cells were similarly incubated with MitoSOX (Invitrogen Cat#M36006) and TMRM (Invitrogen Cat#T668) according to manufacturer’s instructions and acquired live on BD LSR Fortessa analyzer along with propidium iodide (PI). For experiments analyzing ferroptosis, cells were stained with Annexin V for 15 mins at RT in 1x binding buffer (Thermo Fisher Scientific), and fresh propidium iodide (PI) prior to acquisition live on the BD LSR Fortessa analyzer.

<u>Human Tissue</u>: Cells were first incubated with Fixable Blue Live/Dead Dye (Thermo Fisher Scientific) and TruStain FcX (BioLegend) for 15 mins on ice prior to surface staining. Cells were stained for surface markers for 25 mins on ice (antibody details listed in **Supplementary Table 5**). Subsequently for sequencing studies, cells were sorted fresh on BD FACS Aria sorter or Beckman CytoFlex SRT sorter using DAPI or 7-AAD to exclude dead cells. Cells were sorted into pre-chilled FBS-coated microcentrifuge tubes in 200 μl of PBS containing 0.5% BSA.

### In vivo GMP transfer

Bone marrow from indicated donor mice was flushed with cold FACS Buffer using a 27G needle from four long bones (femur and tibia) and filtered through a 70 μm mesh. Samples were subjected to RBC Lysis Buffer (BioLegend) for 2 mins at RT and quenched with ice-cold FACS Buffer prior to downstream processes. Marrow cells were enriched for Kit^+^ progenitors using the Mojosort Mouse Lin-neg enrichment kit (BioLegend #480004). GMPs were sorted into pre-chilled FBS-coated microcentrifuge tubes in 200 μl of PBS containing 0.5% BSA on the Beckman CytoFlex SRT sorter. Cells were spun down and resuspended to concentration of 200K cells/mL in ice-cold PBS. Recipient CD45.1 mice received 100 μl i.v. retro-orbitally under anesthesia at indicated timepoints. Recipient mice were subsequently sacrificed at day 21 and subjected to flow cytometric analysis.

### Mouse CUT&RUN

Low cell input CUT&RUN technique was performed as follows: antibody-stained cell suspensions were lightly fixed in 200 µL of 0.1% formaldehyde (Sigma# 252549) at room temperature for 1 min and then quenched in 10 µL of 2.5M Glycine. 10,000 sorted cells in PBS were mixed with an equal volume of 2X Nuclear Extraction (NE) buffer i.e. 40mM HEPES, 20mM KCl, 0.2% Triton X-100, 40% Glycerol, 2mM DTT, 1mM Spermidine, 2X Roche Complete Protease Inhibitor (Millipore Sigma# 11873580001). 100X of KDAC inhibitor cocktail (100 µM TSA, 50mM sodium butyrate and 50mM nicotinamide in 70% DMSO) was added to the sorted sample for a final concentration of 1X prior to cryopreservation at -80°C. CUT&RUN was performed in collaboration with EpiCypher following a modified CUT&RUN protocol. In brief, samples were thawed and diluted to 1E10^5^ cells/mL in 1X NE buffer. Then, a mixture of 10 µL of activated Concanavalin A (ConA) beads, 2 µL of 1:50 SNAP-CUTANA™ K-MetStat Panel, and 0.5 µg of primary antibody [rabbit IgG (EpiCypher 13-0042; Lot 20335004-04), H3K4me3 (EpiCypher 13-0041; Lot 210760004-01)] was added to 1E10^4^ cells per reaction and incubated overnight. Next day, beads were washed twice with 250 µL Digitonin Buffer [20 mM pH 7.5 HEPES, 150 mM NaCl, 0.5mM Spermidine, 1X Roche Complete mini, 0.01% digitonin] before adding 5 µL of CUTANA pAG-MNase in 50 µL Digitonin Buffer per reaction. Beads were washed twice in Digitonin Buffer and suspended in 50 µL. 2mM CaCl2 was then added to activate MNase and 33 µL High-Salt Stop Buffer [750mM NaCl, 26.4mM EDTA, 5.28mM EGTA, 66 μg/mL RNase A, 66 μg/mL Glycogen] to stop the MNase activity after 2 hr incubation at 4°C. 20 pg of CUTANA E. Coli spike-in DNA was added per sample, followed by a 10 min incubation at 37°C to release the cleaved chromatin. CUT&RUN-enriched DNA were isolated by ConA beads, cleaned up using Serapure beads, and libraries were prepared using a CUTANA CUT&RUN Library Prep Kit (EpiCypher #14-1001). Libraries were pooled and sequenced on Illumina NovaSeq 6000 SP (150- cycle, paired end).

### Single-cell RNA sequencing (scRNA–seq) assay

For each sample, a target recovery of 8000 cells were loaded onto each lane of a 10X Chromium chip according to manufacturer’s instructions. Libraries were prepared according to manufacturer’s instructions. All libraries were quantified via Agilent 2100 hsDNA Bioanalyzer and KAPA library quantification kit (Roche, Cat. #0797014001). Libraries were sequenced at a targeted depth of 25,000 reads per cell using the NovaSeq 6000 S2 100 cycle kit (Illumina).

### Single-cell ATAC sequencing (scATAC–seq) and Multiome

For scATAC-seq preparation, cells were subjected to nuclei isolation following 10x Genomics manufacturer’s protocol with minor adjustments. In case of low-input samples (with <100,000 cells), we utilized 0.2 ml PCR tubes and centrifuged at 4C using swinging rotor buckets to maximize nuclear recovery. For human NSCLC sample multiome assays, cells were subjected to nuclei isolation following 10x Genomics manufacturer’s protocol in the presence of RNAse inhibitor (Sigma) and DTT (Sigma) to prevent RNA degradation. Viability of these nuclei was assessed using Acridine Orange/Propidium Iodide viability staining reagent (Nexcelom), and all samples post-nuclei isolation demonstrated viability at or below 1%. A target recovery number of 8000∼10000 nuclei were loaded onto each lane of a 10X Chromium chip according to manufacturer’s instructions. Barcoded DNA was extracted from the GEMs post-cleanup and amplified with 10x-specific sample indexing following the manufacturer’s protocols. Libraries were quantified using TapeStation (Agilent) and were sequenced in pair-end mode using the NovaSeq 6000 S2 100 cycle kit (Illumina) targeting a depth of 25,000 reads per cell.

### Mouse scRNAseq analysis

Gene expression reads were aligned to the mm10 reference transcriptome and count matrices were generated using the default CellRanger 2.1 workflow, using ‘raw’ matrix output. Following alignment, barcodes matching cells that contained > 500 unique molecular identifiers (UMIs) were extracted. From these cells, those with transcripts >25% mitochondrial genes (QC thresholds min_mc_size = 25, max_f_mit = 0.1) were filtered from downstream analyses. Matrix scaling, logarithmic normalization, and batch correction via data alignment through canonical correlation analysis, and unsupervised clustering using a *K-nn* graph partitioning approach were performed as previously described. Differentially expressed genes were identified using the *FindMarkers* function in Seurat v4.4.0^43^. Alternatively, we used the *metacell* package for clustering cells across the tumor, blood, and bone-marrow samples separately (parameters K=25, T_lfc=3). Subsequently clusters were annotated in a semi-supervised manner using canonical markers for lineage (e.g.- T cell, B cell, Myeloid cell) and myeloid identity clusters were subjected to gene-module analyses as follows: cells were uniformly down-sampled to 2,000 UMI before selecting the set of variable genes. Subsequently, the gene–gene correlation matrix was computed for each sample subsetting for myeloid cells. Correlation matrices were averaged via Fisher Z-transformation. The inverse transformation resulted in the best-estimate correlation coefficients of gene– gene interactions across the dataset. Genes were clustered into modules using complete linkage hierarchical clustering over this correlation distance. Finally, myeloid clusters were annotated using lineage/function-determining gene modules. Where indicated, gene set enrichment analysis was performed (EnrichR) and redundancy-reduced/collapsed results are illustrated (REVIGO). Single-cell gene signature scoring was conducted using UCell v2.4^96^. Analyses were mostly run using mac x86 64- bit platform running R v4.3.1 on macOS Big Sur 11.3.1 or mac x86 64-bit platform running R v4.2.2 on Ubuntu 20.04.4 LTS.

### Mouse scATAC-seq analysis

Fastq files from scATAC-seq samples were aligned to mouse genome reference mm10 using cellranger-atac v2.0.0. Fragment files were parsed with ArchR v1.0.2 and initial quality control (QC) was applied based on sequencing depth and quality (minTSS = 8, minFrags = 5000). Dimensionality reduction and clustering was applied on cells passing filters using standard ArchR workflow^97, 98^. Peaks were called using Macs2^99^ to generate group coverages. To select for myeloid clusters, we relied on ArchR Gene Scores or gene-activity scores (a surrogate for gene expression based on accessibility at the gene loci) of canonical markers and *de novo* marker discovery using *getMarkerFeatures* and *getMarkers* on the GeneScoreMatrix assay (FDR <=0.01 & Log2FC >=1). We filtered out lymphoid cell populations and CD45-negative contaminants, repeating ArchR analysis workflow on these cells to increase the resolution of the monocytic and macrophage compartment. We used *addGeneIntegrationMatrix* with constraints to map between the paired scRNA and scATAC data. Correlation between Gene Scores in scATAC-seq clusters and relative gene expression in scRNA-seq clusters was used to define similarity scores and soft-label annotations to scATACseq clusters in a semi-supervised manner. TF deviations per cluster was inferred following ChromVAR using JAPSAR2020 motifs. Importantly, we calculated differentially accessible marker peak sets for identified monocytic–macrophage clusters and predicted which transcription factors (TFs) mediate binding and define accessibility at these marker peaks using ArchR. Such analysis yields key lineage-determining TFs important for cell identity but can also prioritize TFs that are crucial to that cell’s state and function.

As part of candidate TF nomination, we computed TFs whose gene expression positively correlated with changes in accessibility of their binding motifs (corr >0.4) and ranked by TF deviation (Λ1) across clusters to nominate candidate regulators likely to be important to cellular function and identity. To identify differentially accessible motifs across conditions, we utilized *getMarkerFeatures* on the MotifMatrix assay correcting for TSS enrichment and number of fragments (FDR <=0.05 unless noted). Analyses were run using mac x86 64-bit platform running R v4.3.1 on macOS Big Sur 11.3.1.

For paired analyses of RNA and ATAC features; previously processed and annotated mouse BM, Blood and Tumor scRNAseq Seurat objects were normalized and scaled again using *SCTransform* (glmGamPoi, vars.to.regress = "percent.mt", variable.features.n = 8000). Mouse BM, Blood, and Tumor scATACseq data were processed in parallel as follows; ATAC CellRanger fragment files were transformed into ArrowFiles using ArchR package using above methology. Peaks were called using MACS2 and *addGroupCoverages, addReproduciblePeakSet* and *addPeakMatrix* functions. Peaks co-accessibility matrix was obtained using *addCoAccessibility* function. To pair each scRNAseq cell with a scATACseq cell, datasets from both modalities were split by tissue before running *FindTransferAnchors* and *TransferData* using ATAC cells as reference and RNA cells as query. The combined Seurat object was filtered on myeloid cells (scRNAseq metadata) and previously obtained peak matrix was also subsetted before being incorporated. TF deviations was inferred following ChromVAR using JAPSAR2020 motifs. Downstream gene activation score was calculated for gene sets using UCell v2.4.

For integrated velocity analysis in such paired data, first spliced and unspliced counts loom files were obtained using Velocyto v1.0 from BAM files of respective scRNAseq dataset and loaded in Scanpy v1.9.3. The object was subsetted on myeloid cells based on marker genes and pre-processed through MultiVelo^100^ using top 5000 variable genes. ArchR peak matrix with scATACseq cells renamed with their matched scRNAseq cells was aggregated and feature linkage bedpe file was created based on previously calculated co-accessibility peak matrix using *getCoAccessibility* ArchR function (corCutOff = 0.5, res = 1). MultiVelo *recover_dynamics_chrom* function was used to calculate new velocity that fed CellPath^101^ (num_trajs = 5, flavor = "k-means", num_metacells = 2000). AnnData object was then subsetted on cells belonging to trajectories ending in terminal macrophage clusters. Final velocity and pseudotime were calculated using CellRank^102^ with appropriate kernels.

### Mouse CUT&RUN analysis

FASTQ data were adaptor trimmed using Trim Galore v0.6.6 and aligned to mm10 reference genome using bowtie2 v.2.3.4.3 with parameters -X 700 -I 10. Uniquely mapped, non-duplicated and >MAPQ 30 reads were filtered using SAMToolsview and Picard *MarkDuplicates*. BEDTools intersect was used to remove regions from ENCODE blacklist. Genome coverage tracks were generated using deeptools2 with parameters bamCoverage --binSize 10 –normalizeUsing CPM --extendReads --ignoreDuplicates -- smoothLength 50. Peaks were called using SEACR with parameters ‘norm’ and ‘relaxed’. Paired binary alignment map (BAM) files were generated for each sample after filtering for reads with a mapping quality score > 30 and a sequence length < 150. Bigwig files were derived from fragment files by normalizing read counts, and tracks were visualized using Interactive Genomics Viewer (IGV) v2.17.4. We retrieved a list of myeloid genomic positions associated with open chromatin from the Immunological Genome Project (ImmGen) murine ATAC-seq atlas^103^ to prioritize active or poised states, and subsetted the regions to those retained in the published analysis. We extended the summits ±500 base pairs to form a list of open chromatin regions (OCRs) and trimmed the regions to within chromosomal boundaries using GenomicRanges v1.52.0. These OCRs were then used to derive counts matrices, defined by the number of observed reads in each region, using chromVAR v1.22.1. Counts were then merged per histone mark, empty regions were filtered, and GC bias calculated per region. Using motifmatchr v1.20.0, regions were annotated for presence of TF motifs by referencing a custom database of mouse position frequency matrices. Subsequent chromVAR deviation z-scores indicate the bias-corrected and normalized presence of each histone modification around a particular transcription factor (TF) motif, across all OCRs. To identify differentially variable regions with only 2-3 samples in each condition, we stratified data based on the mean difference between two conditions, and the fraction of intra-sample variance over total variance. The top hits from chromVAR analysis were confirmed by visual inspection on IGV. To unbiasedly identify OCRs which were retained in both tumor-associated GMPs and tumor-associated Ly6Chi monocytes, we applied k-means clustering to chromVAR-corrected OCR scores, and used the web-based tool Genomic Regions Enrichment of Annotations v4.0.4 to annotate and summarize clusters of interest into gene ontology terms. To visualize the underlying distribution of histone modification signal without chromVAR correction, we used deeptools v3.5.5 to generate averaged profiles and heatmaps for each cluster of interest. In brief, bigwigs for each cell type and condition were averaged using *bigwigAverage*, browser extensible data (BED) files were outputted containing regions from each cluster, score matrices were calculated by *computeMatrix*, and plots were generated through *plotHeatmap*.

### Human scATAC-seq and Multiome analysis

Fastq files from scATAC-seq and multiome samples were aligned to human genome reference hg38 using cellranger-atac v2.0.0 or cellranger-arc v2.0.0 respectively. Fragment files were parsed with ArchR v1.0.2 and initial quality control (QC) was applied based on sequencing depth and quality, discarding cells with <3000 fragments or a transcription start site (TSS) enrichment score < 8. Dimensionality reduction and clustering was applied on cells passing filters using ArchR. To select for myeloid clusters, we relied on ArchR Gene Scores of canonical markers and *de novo* marker discovery using *getMarkerFeatures* on the GeneScoreMatrix assay. Thus, we filtered out lymphoid cell populations and CD45negative contaminants for downstream RNA-ATAC integration. snRNA-seq from multiome samples was analyzed using Seurat v4.4.0. Cells with <800 UMI or <400 genes detected were filtered out followed by default functions for data normalization and dimensionality reduction. To aid in the identification of macrophages cell states, we utilized hdWGCNA v0.2.18. This method allows for identification of gene co-expression modules after generation of metacells (groups of neighboring cells in embedding space) to circumvent data sparsity. Metacells were generated with *MetacellsByGroups* function with following parameters: k=20 max_shared=15 and min_cells=20. Gene modules were discovered with function *ConstructNetwork* with following parameters: softpower=3, deepSplit=1, minModuleSize=10, mergeCutHeight=0.2, maxblocksize=35,658. We also verified monocyte and macrophage assignment using orthogonal methodology of consensus non-negative matrix factorization (cNMF; k=55). For each cluster, we annotated using top genes ranked by spectra score in the gene expression program matrix obtained and selected the top 50 genes to generate cellular expression program signatures. In addition to soft assignments, this NMF-based approach can work around challenges of identifying and correcting batch effects emerged during data integration and requires minimal pre-processing steps. To annotate macrophage subtypes in chromatin space; we first co-embedded all scATAC samples and multiome samples together, removed batch effects using harmony v0.1.1. We then annotated each cell cluster by the most abundant macrophage state in Multiome-origin cells that passed QC and were previously annotated in snRNA-seq using marker gene expression. TF activity was inferred using ChromVAR v1.16 implemented by ArchR via *addMotifAnnotations, addBgdPeaks* and *addDeviationsMatrix* functions. For visualization, ArchR built-in plotting functions and ComplexHeatmap v2.10 were used. Subsequently, we carried out similar TF motif enrichment, TF prioritization, and peak2gene linkage analyses as described above in mouse section.

### QUANTIFICATION AND STATISTICAL ANALYSIS

No statistical methods were used *a priori* to determined sample size. Sample size was based on power analyses from prior studies in the lab and upon establishing reproducibility between experiments. For flow cytometry, data was collected on FACSDiva v8 or Beckman CytoExpert, and analyzed using FlowJo v10.9. Absolute cell numbers were calculated using initial loading volume and fluorescent beads (Accucheck Counting Beads PCB100, Molecular Probes) following the manufacturer’s instructions. Where applicable, median fluorescence intensity (MFI) measurements were compared for markers of interest. For histology, scanned H&E slides were quantified by manual annotation of investigator-blinded slides using QuPath v0.4^95^.

Statistical ranges in figures represent individual data points with mean value indicated, unless otherwise indicated in box plots or violin plots where error bars indicate s.e.m. For continuous data satisfying normality assumptions (flow cytometry, histology, and sequencing), statistical significance between conditions was determined using unpaired Student’s t-test for two independent comparisons, or one-way ANOVA with multiple comparison correction for three or more independent groups. Statistical significance between grouped data across three or more conditions was determined using two-way ANOVA with appropriate multiple comparisons’ correction. Statistical significance for mouse survival analyses was performed using Kaplan-Meier log-rank estimate. Statistical analyses for flow cytometry and histology were done using Prism v10.0 (GraphPad). UCell scores calculated on single-cell feature data are based on the Mann-Whitney U statistic and robust to dataset size and sample heterogeneity. DEG analyses in single-cell feature data were subjected to Wilcoxon rank sum test on pseudobulk values as well as ‘bimod’ likelihood-ratio test optimal for single cell feature expression^104^, unless otherwise indicated. Ontology terms in EnrichR were ranked by p-value or odds ratio as indicated; where the enrichment p-value was calculated using Fisher’s exact test comparing the observed frequency with the frequency expected by chance in background gene list. Statistical analyses for sequencing studies were done using native R v4.3.1 function packages.

Notable R packages used: BiocManager v1.30.22; biomaRt v2.56.1; BSgenome v1.68.0; BSgenome.Mmusculus.UCSC.mm10 sequences; BSgenome.Hsapiens.UCSC.hg38 sequences; Seurat v4.9.9; harmony v0.1.1; scDissector v.1.0.0; parallel v4.3.1; ShinyTree v.0.2.7; data.table v.1.14.8; reshape2 v.1.4.4; reticulate v1.32.0; heatmaply v.1.3.0; pheatmap v1.0.12; plotly v.4.10.0; ggvis v.0.4.7; ggplot2 v.3.3.5; cowplot v1.1.1; patchwork v1.1.3; dplyr v.1.3.1; tidyr v1.3.0; tidyverse v2.0.0; plyranges v.1.20.0; Matrix v.0.9.8; seriation v.1.3.5; ArchR v1.0.2; chromVAR v1.22.1; Signac v.1.11.9; complexHeatmap v2.16.0; hdWGCNA v0.2.18; edgeR v3.42.4; limma v3.56.2; presto v1.0.0; EnhancedVolcano v1.18.0; GenomeInfoDb v1.36.1; GenomicRanges v1.52.0; motifmatchr v1.20.0, deeptools v3.5.5, MAST v1.6; Nebulosa v1.10.0; RColorBrewer v1.1; scCustomize v1.1.3; sceasy v.0.0.7; scTools v1.0; SeuratDisk v.0.0.0.9; SeuratWrappers v0.3.19; SignatuR v.0.1.1; SingleCellExperiment v1.22.0; TFBSTools v1.38.0; UCell v2.4.0.

## DATA AND CODE AVAILABILITY

Notable software package versions used in this study are listed. No new software pipelines were used in the study beyond those described in relevant Methods sections. Accession numbers for re-analyzed published datasets are listed in the **Supplementary Table 6**. Processed matrix files and metadata for the mouse tissue scRNA-seq and scATAC-seq generated in this study are made publicly available at the time of publication (**GSE255330**). Processed matrix files and metadata for the human tissue scATAC-seq and 10x multiome data generated in this study are also made publicly available at the time of publication. Any additional information required to interpret data reported in this paper is available from the corresponding author upon reasonable request.

---

**Extended Data Fig. 1:**
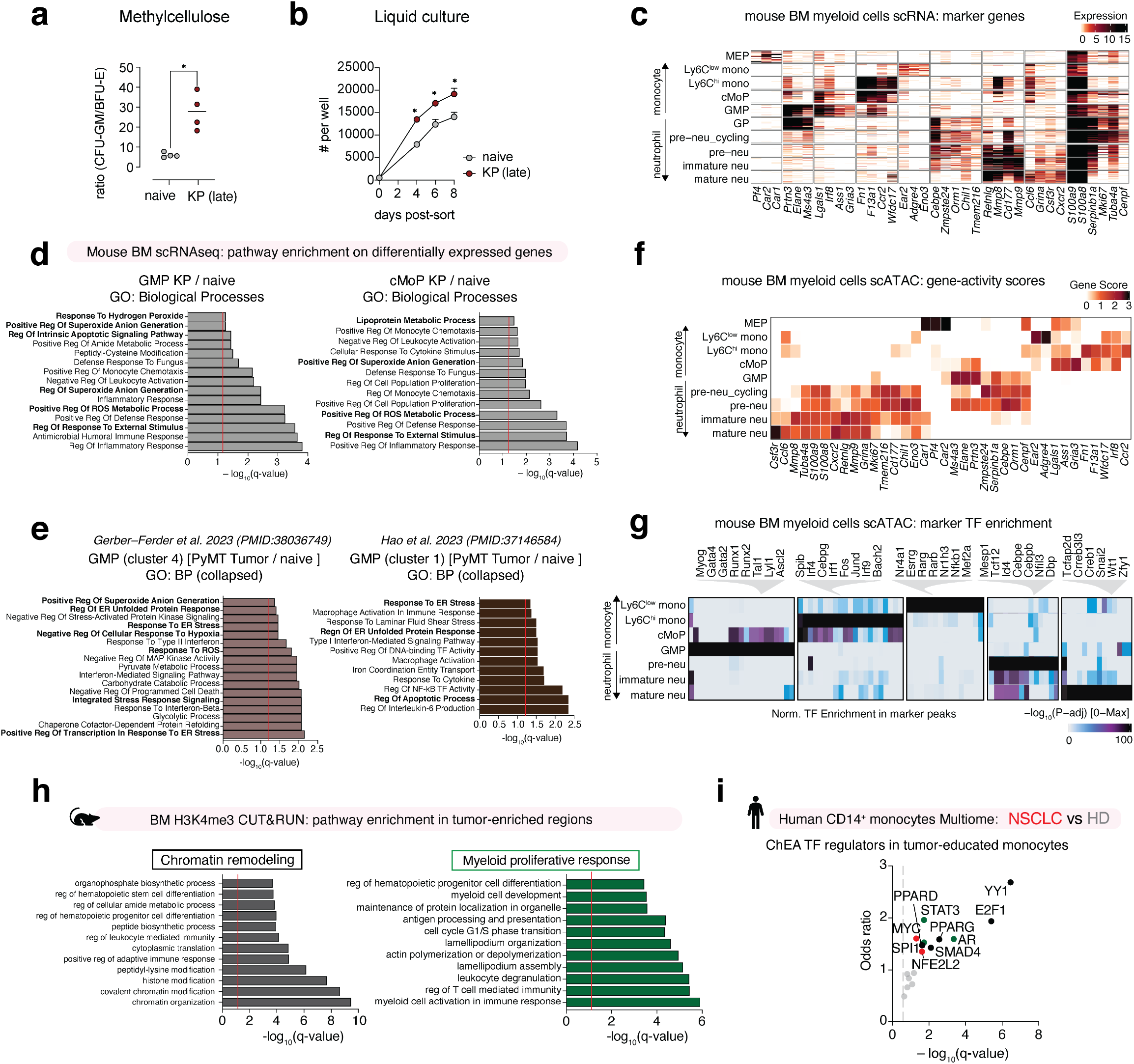
Pathogenic myelopoiesis in cancer associates with changes in the chromatin state of bone marrow myeloid progenitors. a. Ratio of granulocytic-monocytic colony forming units (CFU-GM) to erythroid blasts (BFU-E) formed by BM progenitors from naïve and KP tumor-bearing mice. pooled from 2 independent experiments. N=4 replicates. b. Cell growth quantified during longitudinal liquid culture expansion of sorted BM GMPs from naïve and KP tumor-bearing mice. pooled from 2 independent experiments. N=3 replicates. c. scRNA-seq heatmap of unique molecular identifier (UMI) counts per cell depicting sub-clustering of myeloid cells in BM of naïve and tumor-bearing mice and annotation of cell states based on characteristic markers. Pooled from N=3 mice per group. d. Gene ontology (GO): Biological process (BP) terms enriched in KP tumor-bearing mouse BM GMPs (LEFT) and cMoPs (RIGHT) compared to naïve counterpart. Curated terms arranged by adjusted p-value (log q-value). *p*-values computed by hypergeometric test with multiple test correction. e. Gene ontology (GO): Biological process (BP) terms enriched in PyMT tumor-bearing mouse GMPs compared to naïve counterpart; data obtained from Gerber-Ferder et al. 2023 (LEFT) and Hao et. al. 2023 (RIGHT). Curated terms arranged by adjusted p-value (log q-value). *p*-values computed by hypergeometric test with multiple test correction. f. scATAC-seq heatmap depicting column-normalized gene scores across indicated myeloid cell states in BM of tumor-bearing and naïve mice. Pooled from N=3 mice per group. g. scATAC-seq heatmap depicting normalized score for transcription factor (TF) motif accessibility enriched in marker peak regions of indicated myeloid cell states in BM of naïve and tumor-bearing mice. Pooled from N=3 mice per group. h. Pathways in indicated clusters of H3K4me3 peaks enriched in BM GMPs and Ly6C^hi^ monocytes from KP tumor-bearing mice compared to naïve mice. Curated terms arranged by adjusted p-value (log q-value). i. ChIP-X Enrichment Analysis (ChEA) calculated TF regulators on differentially expressed genes in CD14 monocytes from blood of patients with NSCLC compared to healthy donors. TFs arranged by adjusted p-value (log q-value). Dot color indicating major known biological pathways. *p*-values computed by Welch’s t-test (a). *p*-values computed by multiple unpaired t-test across timepoints (b). *p*-values computed by hypergeometric test with multiple test correction (d),(e),(h),(i). P-value of < 0.05 denoted *; p-values < 0.01 denoted **; p-values < 0.001 denoted ***.

**Extended Data Fig. 2:**
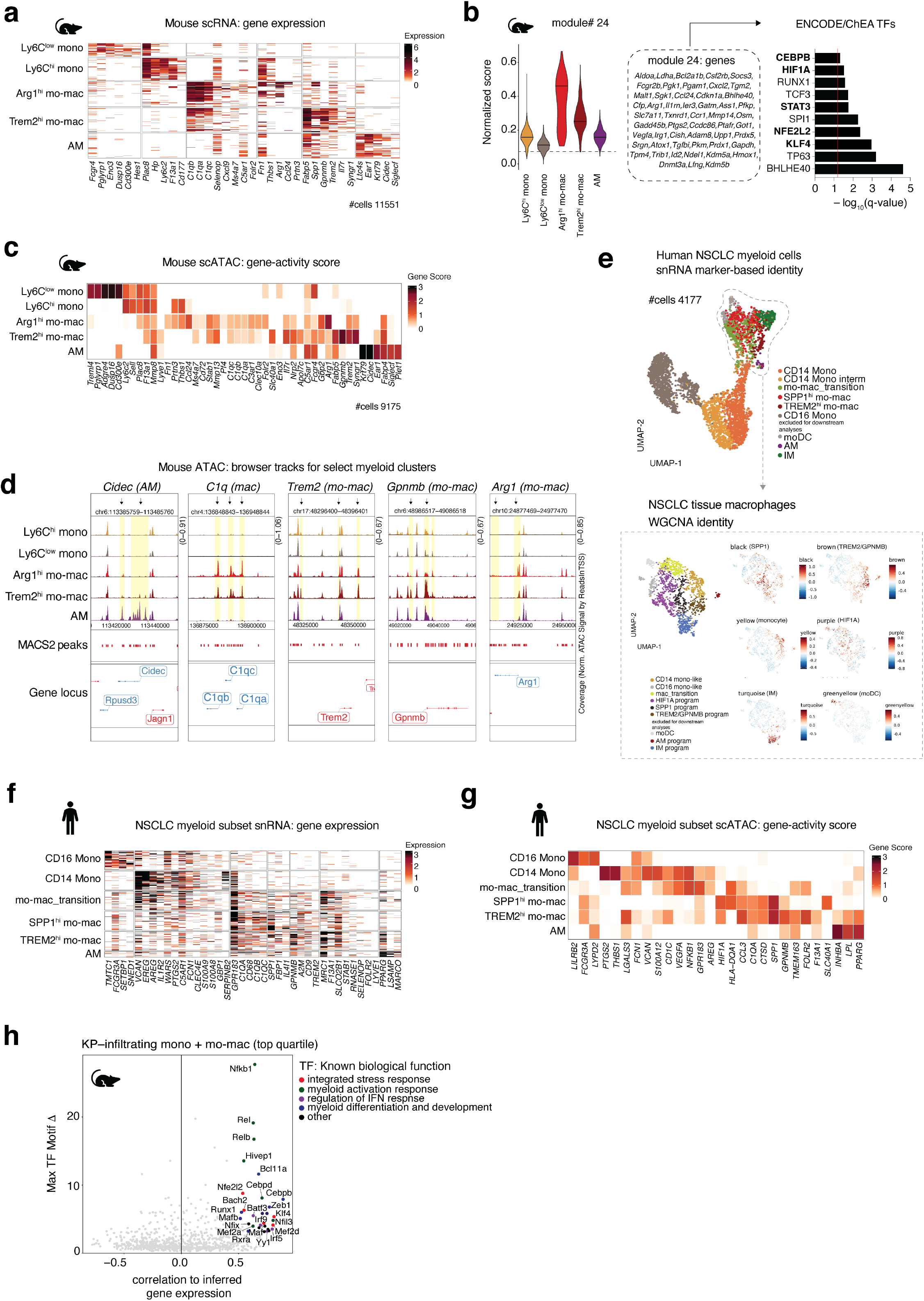
Tumor myelopoiesis fuels mo-macs in TME with sustained cytoprotective stress responses. a. scRNA-seq heatmap plot of UMI counts per cell depicting sub-clustering of myeloid cells in lung of naïve and KP tumor-bearing mice and annotation of cell states based on characteristic markers. N=3 mice pooled for one experiment. b. Aggregate module score for metabolic and cytoprotective gene program across KP lung tumor-infiltrating myeloid cell clusters (LEFT), with ENCODE/ChEA3 calculated TF regulators on identified gene module. c. scATAC-seq heatmap depicting column-normalized gene scores across indicated myeloid cell states in lung of naïve and tumor-bearing mice. N=4 pooled for one experiment. d. Representative browser track plots at known marker gene loci for myeloid cell states identified in scATAC-seq data from lung of naïve and KP tumor-bearing mice. Yellow highlights and arrows indicate peak regions of interest. e. UMAP depicting myeloid cells in snRNA-seq data from human NSCLC primary lung tumors (TOP; N=5 patients); colored by marker gene-based cellular identity. Representative UMAP depicting WCGNA-based identity for specific macrophage gene programs in subsetted dataset with exemplar weighted programs (BOTTOM). f. snRNA-seq heatmap of UMI counts per cell depicting sub-clustering of myeloid cells in human NSCLC primary lung tumors and annotation based on characteristic markers. N=5 patients pooled. g. scATAC-seq heatmap depicting column-normalized gene scores across indicated myeloid cell states in lung tumors of patients with NSCLC. N=14 patients pooled. h. scATAC-seq candidate TF regulators in mouse tumor-infiltrating monocyte and macrophage clusters, prioritized by maximum TF motif deviation (Δ) across clusters. Dot color indicating major known biological pathway. N=4 pooled. *p*-values computed by hypergeometric test with multiple test correction (b).

**Extended Data Fig. 3:**
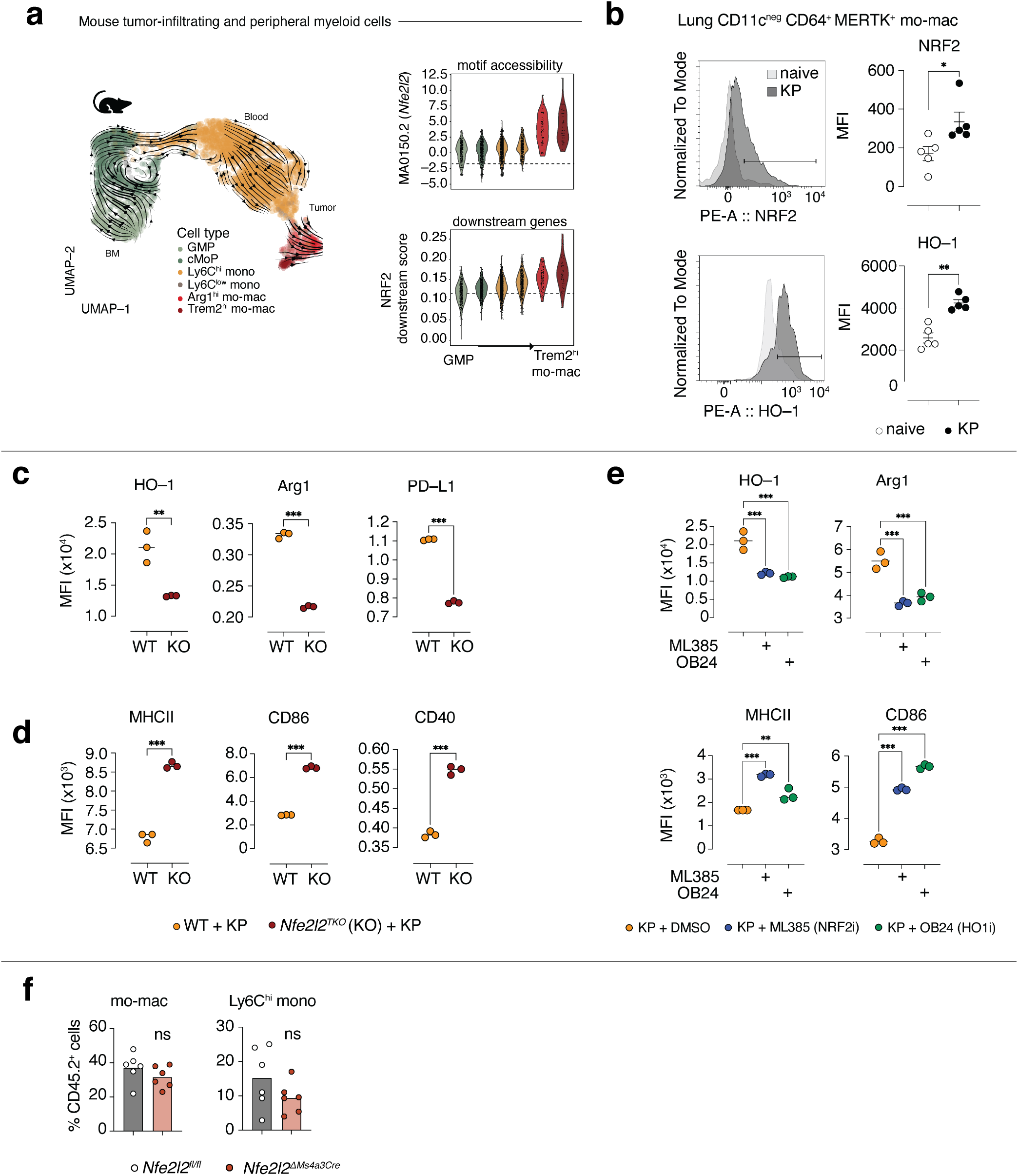
Stepwise activation of NRF2 signaling regulates tumor-associated mo-mac survival and immunosuppressive function. a. UMAP plot of paired mouse scRNA-seq labeled by cell type identity (LEFT) with quantification of *Nfe2l2* TF motif accessibility and aggregate score for NRF2 downstream gene program activation (RIGHT). b. Representative flow cytometry histogram and median fluorescence intensity (MFI) quantification of NRF2 and HO-1 expression in lung-infiltrating CD64^+^MERTK^+^ mo-macs of naïve or KP tumor-bearing mice. N=5-6 mice per group. c. Relative MFI quantification of HO-1, immunoregulatory markers Arg1, PDL1, CD206 in NRF2KO (KO) or control (WT) bone-marrow derived macrophages exposed to KP tumor conditioned media. N=3 per group. d. Relative MFI quantification of immunostimulatory markers MHCII, CD80, CD86, and CD40 in NRF2KO (KO) or control (WT) bone-marrow derived macrophages exposed to KP tumor conditioned media. N=3 per group. e. Relative MFI quantification of HO-1, Arg1, MHCII, CD86 in KP conditioned media–exposed BMDMs treated with ML385 (NRF2 inhibitor) and OB24 (HO–1 inhibitor). N=3 per group. f. *In vivo* tracing of KP tumor-primed BM GMPs transferred from Nfe2l2^ΔMs4a3^ mice or Nfe2l2^fl/fl^ control littermates into KP tumor-bearing congenic CD45.1 hosts, with relative abundance of donor-derived tumor mo-macs and Ly6C^hi^ monocytes. N=6 per group. One experiment. *p*-values computed by one-way ANOVA with Dunnett’s multiple comparisons test (e). *p*-values computed by unpaired t-test (b),(c),(d),(f). P-value of < 0.05 denoted *; p-values < 0.01 denoted **; p-values < 0.001 denoted ***.

**Extended Data Fig. 4:**
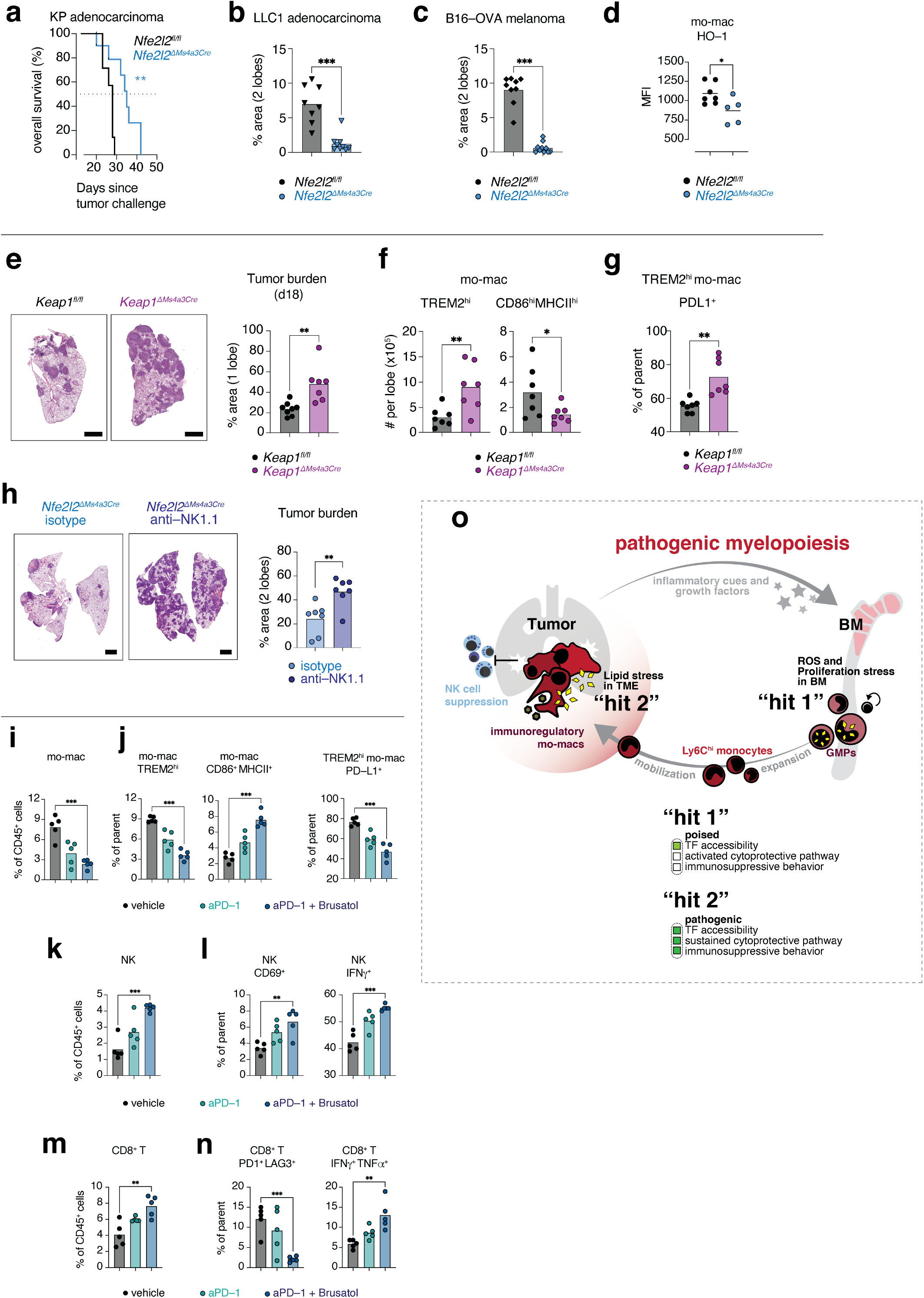
NRF2 signaling sustains myelopoiesis promoting NK and T cell immunosuppression in TME. a. Kaplan-Meier plot depicting overall survival of KP tumor-bearing Nfe2l2^ΔMs4a3^ mice or Nfe2l2^fl/fl^ negative littermates. N=8–10 mice per group. b. Representative lungs from LLC1 tumor-bearing Nfe2l2^ΔMs4a3^ mice or Nfe2l2^fl/fl^ negative littermates with quantification of tumor burden. N=9–10 mice per group. Data are individual data points with bar denoting mean. c. Representative lungs from B16-OVA metastatic tumor-bearing Nfe2l2^ΔMs4a3^ mice or Nfe2l2^fl/fl^ negative littermates with quantification of tumor burden. N=9–10 mice per group. d. MFI quantification of HO-1 expression in lung-infiltrating mo-macs of Nfe2l2^ΔMs4a3^ mice or Nfe2l2^fl/fl^ littermates. N=7–8 mice per group. e. Representative lungs from KP tumor-bearing Keap1^ΔMs4a3^ mice or Keap1^fl/fl^ littermates with quantification of tumor burden. N=7–8 mice per group. f. Number of lung-infiltrating mo-macs expressing GPNMB and CD9 (TREM2^hi^), CD86 and MHCII (CD86^+^MHCII^+^) in KP tumor-bearing Keap1^ΔMs4a3^ mice or Keap1^/fl^ negative littermates. N=7–9 mice per group. g. Frequency of lung-infiltrating TREM2^hi^ mo-macs expressing immunoregulatory marker PDL1 in KP tumor-bearing Keap1^ΔMs4a3^ mice or Keap1^fl/fl^ negative littermates. N=6 mice per group. h. Representative histology at day 21 from KP tumor-bearing Nfe2l2^ΔMs4a3^ mice that received either anti-NK1.1 depletion antibodies or isotype, with quantification of tumor burden. N=7–9 mice per group. i. Frequency of mo-macs in lung tumors of tumor-bearing mice treated as indicated. N=5–7 mice per group. j. Frequency of mo-macs expressing GPNMB and CD9 (TREM2^hi^), CD86 and MHCII (CD86^+^MHCII^+^) and frequency of TREM2^hi^ mo-macs expressing PDL1 in tumor-bearing mice treated as indicated. N=5–7 mice per group. k. Frequency of NK cells in tumor-bearing mice treated as indicated. N=5–7 mice per group. l. Percentage of NK cells expressing activating marker CD69 and producing IFNγ in tumor-bearing mice treated as indicated. N=5–7 mice per group. m. Frequency of CD8^+^ T cells in tumor-bearing mice treated as indicated. N=5–7 mice per group. n. Frequency of CD8^+^ T cells expressing inhibitory markers PD1, LAG3 and producing IFNγ and TNFα in tumor-bearing mice treated as indicated. N=5-7 mice per group. o. Model depicting “two subsequent hits” within the myeloid lineage during tumor-induced myelopoiesis that promote myeloid suppression in the TME. The first hit is in the BM where ROS and proliferative stress in response to tumor-induced expansion (1) poises NRF2 programs in myeloid progenitors and (2) initiates the pathogenic differentiation of Ly6C^hi^ monocytes. This is then compounded by a second hit which occurs when circulating Ly6C^hi^ monocytes accumulate within the lipid- and ROS-laden TME (3) which further solidifies NRF2 program to promote differentiation into immunosuppressive long-lived mo-macs (4). p-values computed by Log-rank (Mantel-Cox) test (a). *p*-values computed by unpaired t-test (b)–(h). *p*-values computed by one-way ANOVA with Dunnett’s multiple comparisons test (i)–(n). P-value of < 0.05 denoted *; p-values < 0.01 denoted **; p- values < 0.001 denoted ***.s

