## Supplementary Table 5 for "Myeloid progenitor dysregulation fuels immunosuppressive macrophages in tumors"

**Supplementary Table 5: Antibodies used for Mouse and Human flow cytometry**

| Reagent or Resource | Source | Cat# | Identifier |
|---|---|---|---|
| **Antibodies Mouse (clone)** | | | |
| anti-mouse CD45 (30-F11) | Biolegend | 103138 | RRID:AB_2563061 |
| anti-mouse CD62L (MEL-14) | Biolegend | 104406 | RRID:AB_313093 |
| anti-mouse CXCR4 (2B11) | Thermo Fisher Scientific | 12-9991-82 | RRID:AB_891391 |
| anti-mouse CD88 (20/70) | Biolegend | 135813 | RRID:AB_2750209 |
| anti-mouse CD117 (2B8) | Thermo Fisher Scientific | 25-1171-82 | RRID:AB_469644 |
| anti-mouse Ly6A/E (D7) | Thermo Fisher Scientific | 56-5981-82 | RRID:AB_657836 |
| anti-mouse CD11b (M1/70) | Thermo Fisher Scientific | 48-0112-82 | RRID:AB_1582236 |
| anti-mouse MHCII I-A/I-E (M5/114.15.2) | Biolegend | 107636 | RRID:AB_2561397 |
| anti-mouse Ly6C (HK1.4) | Biolegend | 128035 | RRID:AB_2562353 |
| anti-mouse CD115 (AFS98) | Thermo Fisher Scientific | 13-1152-85 | RRID:AB_466565 |
| anti-mouse Siglec-F (1RNM44N) | Thermo Fisher Scientific | 78-1702-80 | RRID:AB_2744907 |
| anti-mouse CD34 (RAM34) | Thermo Fisher Scientific | 11-0341-82 | RRID:AB_465022 |
| anti-mouse CD48 (HM48-1) | Thermo Fisher Scientific | 46-0481-80 | RRID:AB_10870793 |
| anti-mouse CD135 (A2F10) | Biolegend | 135306 | RRID:AB_1877217 |
| anti-mouse CD150 (TC15-12F12.2) | Biolegend | 115924 | RRID:AB_2270307 |
| anti-mouse CD16/32 (93) | Biolegend | 553143 | RRID:AB_394658 |
| anti-mouse CD16/32 (2.4G2) | Thermo Fisher Scientific | 56-0161-82 | RRID: AB_394656 |
| anti-mouse CD11c (N418) | Thermo Fisher Scientific | 47-0114-82 | RRID:AB_1548652 |
| anti-mouse CD11b (M1/70) | Thermo Fisher Scientific | 47-0112-82 | RRID:AB_1603193 |
| anti-mouse CD44 (IM7) | Biolegend | 103041 | RRID:AB_2571953 |
| anti-mouse Ly6G (1A8) | Biolegend | 127624 | RRID:AB_10645331 |
| anti-mouse TER119 (TER-119) | Thermo Fisher Scientific | 47-5921-82 | RRID:AB_1548786 |
| anti-mouse NK1.1 (PK136) | Biolegend | 108724 | RRID:AB_830870 |
| anti-mouse CD3e (17A2) | Thermo Fisher Scientific | 47-0032-82 | RRID:AB_1272181 |
| anti-mouse B220 (RA3-6B2) | Thermo Fisher Scientific | 47-0452-82 | RRID:AB_1518810 |
| anti-mouse CD25 (PC61) | Biolegend | 102028 | RRID:AB_2295974 |
| anti-mouse CD335 (29A1.4) | Thermo Fisher Scientific | 12-3351-82 | RRID:AB_1210743 |
| anti-mouse PD-1 (J43) | Thermo Fisher Scientific | 25-9985-82 | RRID:AB_10853805 |
| anti-mouse LAG-3 (C9B7W) | Biolegend | 125209 | RRID:AB_1063972 |
| anti-mouse TIM-3 (8B.2C12) | Thermo Fisher Scientific | 17-5871-82 | RRID:AB_2573234 |
| anti-mouse CD4 (GK1.5) | Biolegend | 100430 | RRID:AB_493699 |
| anti-mouse CD8a (53-6.7) | Biolegend | 100750 | RRID:AB_2562610 |
| anti-mouse CD3e (eBio500A2) | Thermo Fisher Scientific | 48-0033-82 | RRID:AB_2016704 |
| anti-mouse KLRG1 (2F1/KLRG1) | Biolegend | 138405 | RRID:AB_10578565 |
| anti-mouse PD-L1 (MIH5) | Thermo Fisher Scientific | 12-5982-82 | RRID:AB_466089 |
| anti-mouse PD-L2 (TY25) | Thermo Fisher Scientific | 12-5986-82 | RRID:AB_466097 |
| anti-mouse CD40 (1C10) | Thermo Fisher Scientific | 17-0401-82 | RRID:AB_469386 |
| anti-mouse CD86 (GL-1) | Biolegend | 105022 | RRID:AB_493466 |
| anti-mouse CD80 (16-10A1) | Biolegend | 104725 | RRID:AB_10900989 |
| anti-mouse MERTK (2B10C42) | Biolegend | 151503 | RRID:AB_2617035 |
| anti-mouse F4/80 (BM8) | Biolegend | 123106 | RRID:AB_893501 |
| anti-mouse CD64 (X54-5/7.1) | Biolegend | 139318 | RRID:AB_2566557 |
| anti-mouse CD206 (C068C2) | Biolegend | 141720 | RRID:AB_2562248 |
| anti-mouse CD11c (N418) | Biolegend | 117336 | RRID:AB_2565268 |
| anti-mouse CD2 (RM2-5) | Biolegend | 100114 | RRID:AB_2563092 |
| anti-mouse CD19 (eBio1D3) | Thermo Fisher Scientific | 48-0193-82 | RRID:AB_2734905 |
| anti-mouse CD45.2 (104) | Thermo Fisher Scientific | 45-0454-82 | RRID:AB_953590 |
| anti-mouse CD45.1 (A20) | Biolegend | 110741 | RRID:AB_2563378 |
| anti-mouse NKG2D (CX5) | Thermo Fisher Scientific | 13-5882-85 | RRID:AB_466746 |
| anti-mouse CD301/CLEC10A (LOM-14) | Biolegend | 145707 | RRID:AB_2562942 |
| anti-mouse XCR1 (ZET) | Biolegend | 148220 | RRID:AB_2566410 |
| anti-mouse LYVE-1 (ALY7) | Thermo Fisher Scientific | 53-0443-80 | RRID:AB_1633417 |
| anti-mouse IFNg (XMG1.2) | Biolegend | 505850 | RRID:AB_2616698 |
| anti-mouse TNFa (MP6-XT22) | Biolegend | 506307 | RRID:AB_315429 |
| anti-mouse FOXP3 (FJK-16s) | Thermo Fisher Scientific | 17-5773-82 | RRID:AB_469457 |
| anti-mouse CD157 (BP-3) | Biolegend | 140207 | RRID:AB_10901172 |
| anti-mouse CD319 (4G2) | Biolegend | 152005 | RRID:AB_2632677 |
| anti-mouse Ly6G (1A8) | Biolegend | 127645 | RRID:AB_2566317 |
| anti-mouse CD177 | R&D Systems | FAB8186P025 | N/A |
| anti-mouse Arg1 (A1exF5) | Thermo Fisher Scientific | 48-3697-82 | RRID:AB_2734837 |
| anti-mouse Relma (DSBRELM) | Peprotech | 500-P214BT | RRID:AB_1268707 |
| anti-mouse CD163 (S15049I) | Biolegend | 155309 | RRID:AB_2814063 |
| anti-mouse GPNMB (CTSREVL) | Thermo Fisher Scientific | 50-5708-82 | RRID:AB_2574239 |
| anti-mouse CD9 (MZ3) | Biolegend | 124817 | RRID:AB_2783077 |
| anti-mouse ki67 (SolA15) | Thermo Fisher Scientific | 11-5698-82 | RRID:AB_11151330 |
| anti-mouse/human HO-1 (HO-1-2) | Enzo | ENZ-ABS687-0200 | N/A |
| anti-mouse/human HO-1 (HO-1-2) | Enzo | ADI-OSA-111-F | RRID:AB_10618556 |
| Streptavidin BV650 | Biolegend | 405232 | N/A |
| Streptavidin BV510 | Biolegend | 405233 | N/A |
| Streptavidin APC-e780 | Thermo Fisher Scientific | 47-4317-82 | RRID:AB_10366688 |
| anti-mouse NK1.1 (PK136) | BioXCell | #BE0036 | RRID:AB_1107737 |
| anti-mouse CD8a (2.43) | BioXCell | #BE0061 | RRID: AB_1125541 |
| anti-mouse IgG2a (2A3) | BioXCell | #BE0085 | RRID: AB_1107769 |
| anti-mouse IgG2b (LTF-2) | BioXCell | #BE0090 | RRID: AB_1107780 |
| anti-mouse PD-1 (RMP1-14) | BioXCell | #BE0146 | RRID:AB_10949053 |
| anti-mouse PD-L1 (10F.9G2) | BioXCell | #BE0101 | RRID:AB_10949073 |
| **Antibodies Human (clone)** | | | |
| Human TruStain FcX | Biolegend | 422302 | RRID:AB_2818986 |
| anti-human CD34 (AC136) | Miltenyi Biotec | 130-113-178 | RRID:AB_2726005 |
| anti-human HLA-DR | BD Biosciences | 552764 | RRID:AB_394453 |
| anti-human CD14 (M5E2) | Biolegend | 301839 | RRID:AB_2563425 |
| anti-human CD38 (HIT2) | Biolegend | 303515 | RRID:AB_2072782 |
| anti-human CD90 (5E10) | Biolegend | 328119 | RRID:AB_2203302 |
| anti-human CD45RA (HI100) | Biolegend | 304127 | RRID:AB_10708880 |
| anti-human CD49f (GoH3) | Biolegend | 313619 | RRID:AB_2128022 |
| anti-human CD16 (3G8) | Biolegend | 302017 | RRID:AB_314218 |
| anti-human CD66b (G10F5) | Biolegend | 305114 | RRID:AB_2566038 |
| anti-human CD11c (3.9) | Biolegend | 301612 | RRID:AB_493021 |
| anti-human CD20 (2H7) | Biolegend | 302349 | RRID:AB_2565524 |
| anti-human CD3 (5K7) | Biolegend | 344820 | RRID:AB_10662538 |
| anti-human CD56 (5.1H11) | Biolegend | 362535 | RRID:AB_2565652 |
| Streptavidin BV605 | Biolegend | 405229 | N/A |
