## Supplementary Table 6 for "Myeloid progenitor dysregulation fuels immunosuppressive macrophages in tumors"

**Supplementary Table 6: Accession codes**

| Software and algorithms | | | |
|---|---|---|---|
| **Resource** | **Source** | **URL** | **Identifier** |
| Flowjo v10.5 | Flowjo, L.L.C. | N/A | RRID: SCR_008520 |
| Prism v9.0 | Graphpad | N/A | RRID: SCR_002798 |
| FIJI v | ImageJ | N/A | RRID: SCR_002285 |
| QuPath v0.4 | QuPath | N/A | RRID:SCR_018257 |
| Cell Ranger | 10x Genomics | N/A | RRID:SCR_023221 |
| Cell Ranger ATAC | 10x Genomics | N/A | RRID:SCR_023221 |
| Cell Ranger ARC | 10x Genomics | N/A | RRID:SCR_023221 |
| R v4.3.1 | R Foundation | N/A | RRID:SCR_001905 |
| Rstudio v12.1 | Posit | N/A | RRID:SCR_000432 |
| Python 3.8.18 | Python | N/A | RRID:SCR_008394 |
| ArchR v1.0.2 | github.com/GreenleafLab/ArchR | github.com/GreenleafLab/ArchR | RRID:SCR_020982 |
| Macs2 | github.com/macs3-project/MACS | github.com/macs3-project/MACS | RRID:SCR_013291 |
| Seurat v4.4.0 | github.com/satijalab/seurat | github.com/satijalab/seurat | RRID:SCR_016341 |
| scDissector | github.com/effiken/scDissector | github.com/effiken/scDissector | N/A |
| UCell v2.4 | github.com/carmonalab/UCell | github.com/carmonalab/UCell | N/A |
| pycistopic v1.0.3 | github.com/aertslab/pycisTopic | github.com/aertslab/pycisTopic | N/A |
| scanpy v1.9.3 | pypi.org/project/scanpy/ | pypi.org/project/scanpy/ | N/A |
| pycistarget v1.0.3 | github.com/aertslab/pycistarget | github.com/aertslab/pycistarget | N/A |
| velocyto v1.0 | github.com/velocyto-team/velocyto.py | github.com/velocyto-team/velocyto.py | N/A |
| MultiVelo | github.com/welch-lab/MultiVelo | github.com/welch-lab/MultiVelo | N/A |
| CellPath v0.1.0 | github.com/PeterZZQ/CellPath | github.com/PeterZZQ/CellPath | N/A |
| CellRank v2.0.2 | github.com/theislab/cellrank | github.com/theislab/cellrank | N/A |
| **Deposited Data** | | | |
| PyMT and naïve BM HSPCs scRNA-seq | Hao et al 2023 | doi.org/10.1016/j.stem.2023.04.005 | GSE188648 |
| PyMT and naïve BM progenitor scRNA-seq | Gerber-Ferder et al 2023 | doi.org/10.1038/s41556-023-01291-w | GSE243964 |
| KP and naïve BM myeloid progenitor scRNA-seq | LaMarche et al 2024 | doi.org/10.1038/s41586-023-06797-9 | GSE245236 |
| KP and naïve BM myeloid progenitor H3K4me3 CUT&RUN | ImmGen (in preparation) | rstats.immgen.org/cutrun/index.html | N/A |
| KP and naïve BM myeloid progenitor scATAC-seq | This paper | N/A | GSE255330 |
| KP and naïve Blood myeloid scRNA-seq | This paper | N/A | GSE255330 |
| KP and naïve Blood myeloid scATAC-seq | This paper | N/A | GSE255330 |
| KP and naïve tissue myeloid scRNA-seq | This paper | N/A | GSE255330 |
| KP and naïve tissue myeloid scATAC-seq | This paper | N/A | GSE255330 |
| Human Lung cancer scRNA-seq (multiome) | This paper | N/A | GSE255330 |
| Human Lung cancer scATAC-seq (multiome) | This paper | N/A | GSE255330 |
| Human Lung cancer PBMC scRNA-seq | This paper | N/A | GSE255330 |
| Human Lung cancer PBMC scRNA-seq | This paper | N/A | GSE255330 |
| Human Lung cancer scRNA-seq | Leader et al 2021 | doi.org/10.1016/j.ccell.2021.10.009 | GSE154826 |
